# The cold-inducible RNA-binding protein RBM3 stabilises viral mRNA at cooler temperatures representative of the upper respiratory tract

**DOI:** 10.1101/2024.07.25.605104

**Authors:** Hannah L Coutts, Swathi Sukumar, Hannah L Turkington, Stefano Bonazza, Emily Peate, Erin M P E Getty, Ultan F Power, David G Courtney

**Author notes:** These authors contributed equally (Hannah L Coutts, Swathi Sukumar).

## Abstract

Temperature is a critical determinant of host-pathogen interactions in the respiratory tract, where influenza A virus (IAV) has adapted to the cooler environment of the upper respiratory tract (URT) to enable efficient replication and transmission. The cold-inducible RNA-binding protein RBM3 is highly expressed in nasopharyngeal tissue and is known to stabilise mRNAs under hypothermic conditions; however, its role during viral infection has not been defined.

Here, we identify RBM3 as a key host factor that facilitates IAV replication at sub-physiological temperatures. siRNA-mediated knockdown of RBM3 significantly impaired viral replication, while its constitutive overexpression at 37°C restored replication to levels typically observed at 33°C. Mechanistically, RBM3 binds directly to viral nucleoprotein (NP) mRNA, prolonging mRNA half-life and inevitably leading to increased viral production. This effect was abolished in an RNA-binding-deficient RBM3 mutant, confirming the requirement for direct RNA interaction, visualised by smiFISH-PLA. Crucially, this positive regulation of NP mRNA and interaction was validated in well-differentiated primary nasal epithelial cells, highlighting the physiological relevance of RBM3 in the human URT.

These results reveal a temperature-sensitive host-virus interaction that promotes IAV replication in the cooler URT, a key site for viral shedding and transmission. By linking environmental temperature, host RNA-binding proteins, and viral mRNA stability, this study uncovers a novel mechanism of respiratory virus adaptation and identifies RBM3 as a potential therapeutic target for limiting early-stage viral replication and transmission.

**Author Summary:** To establish productive infection and ensure transmission, respiratory viruses such as influenza A virus (IAV) must adapt to the diverse environments of the human respiratory tract. One key challenge is the temperature gradient that exists between the upper and lower respiratory tract. The upper respiratory tract (URT), the main site of viral entry and transmission, maintains a lower physiological temperature (∼33°C) than the lower respiratory tract (∼37°C, LRT), creating a distinct cellular environment. In this study, we investigated how this cooler URT temperature alters the landscape of RNA-binding proteins (RBPs), which play a central role in regulating gene expression after transcription. Using mass spectrometry-based profiling and molecular virology approaches, we identified a cold-inducible RBP, RBM3, as significantly enriched at 33°C and acting as a key proviral host factor during IAV infection. We show that RBM3 binds directly to viral nucleoprotein (NP) mRNA, mainly in the cytoplasm, in both immortalised cell models and primary nasal epithelial cells. This interaction promotes the stability of NP mRNA thereby enhancing the overall production of infectious virions. Importantly, disrupting RBM3’s RNA-binding ability abolished this proviral effect. These findings reveal that RBM3, elevated in the cooler URT environment, directly supports IAV replication. This work highlights how subtle physiological differences in host tissue can reshape post-transcriptional regulation and influence the outcome of respiratory virus infection.

## Introduction

Respiratory viruses remain an ever-present pandemic concern, in part due to their efficient airborne mode of transmission [1]. While severe pathology from respiratory viruses such as influenza A virus (IAV) or SARS-CoV-2 is typically associated with replication in the lower respiratory tract (LRT), transmission predominantly occurs via small droplets and aerosols expelled from the upper respiratory tract (URT) [1–4]. Thus, advancing research into the unique cellular environment of the nasopharyngeal epithelium is critical to deepening our understanding of respiratory virus transmission [5] and will ultimately support public health efforts to develop the next generation of nasal vaccines that boost mucosal immunity [6,7].

A key feature of this URT environment is its lower physiological temperature, with nasopharyngeal epithelial cells typically residing at or below 33°C, several degrees cooler than the average core body temperature of 37°C [8]. Though seemingly minor, this temperature difference has far-reaching effects on cellular processes, including viral replication kinetics [9]. Because molecular interactions and reaction rates are governed by thermodynamic principles, temperature is a fundamental variable in regulating biochemical activity. Accordingly, the interaction landscape, including RNA and protein folding, stability, and binding, is substantially altered under cooler conditions [10]. These changes can lead to RNA or protein misfolding [10–12], and, crucially, to temperature-sensitive differences in RNA-binding protein (RBP) affinities for target RNAs, with wide-ranging effects on post-transcriptional regulation [13].

Temperature-dependent variation in RBP-RNA binding has significant implications for mRNA spatial localisation, stability, and translation, processes that can influence viral RNA processing and immune evasion [14]. While prior studies have observed temperature-driven changes in transcript and protein abundance [15,16], it remains poorly understood how growth temperature alters the global landscape of RBPs that bind to cellular mRNAs—collectively known as the RBPome.

The central aim of this study was to characterise the post-transcriptional environment encountered by respiratory viruses, with a focus on IAV, and to determine how altered growth temperature affects viral mRNA regulation. Specifically, we profiled the mRNA-bound RBPome of lung epithelial cells (A549 cells) grown at 37°C and 33°C, mimicking LRT and URT temperatures, respectively. Recent studies have highlighted the critical role of the host RBPome in shaping viral replication by regulating nuclear export [17], translation [18], and genome replication [19], underscoring the importance of this approach [18,20].

Our data reveal extensive temperature-dependent variation in RBPs associated with cellular mRNAs, identifying large sets of RBPs whose binding is either enriched or depleted at 33°C relative to 37°C. Among the most striking findings was the cold-induced enrichment of RNA Binding Motif 3 (RBM3), a well-characterised cold-shock protein known to support cell survival under hypothermic stress [21,22]. Notably, RBM3 was recently identified by us and our collaborators in an IAV NP mRNA-specific interactome screen [23]. Here, we build on that work and demonstrate that in A549 cells RBM3 functions as a proviral host factor during IAV infection by directly binding & stabilising viral NP mRNA transcripts.

Crucially, we validate these findings in well-differentiated primary nasal epithelial cells (PNECs) derived from multiple human donors through a combination of smiFISH-PLA and antisense oligonucleotide (ASO)-mediated induction of RBM3 expression [24] followed by infection. Notably, by using our novel smiFISH-PLA technique in both immortalised A549s and PNECs, we visualise the direct interaction between a viral transcript and a host RBP for the first time.

Together, these findings reveal that RNA regulation in the URT occurs within a distinct post-transcriptional landscape, shaped by cooler tissue temperature. This environment profoundly impacts viral replication dynamics and represents a key barrier that respiratory viruses, such as IAV, have overcome to establish infection and achieve transmission. Understanding these adaptations not only provides new insights into viral evolution and host-pathogen interactions but also offers a foundation for developing temperature-sensitive antiviral strategies and next-generation respiratory vaccines.

## Results

### The cold RBPome of A549 cells reveals RBM3 as a significantly enriched mRNA interactor

As previously mentioned, the temperature gradient normally observed in the respiratory tract is known to yield distinct variations in cellular transcriptomes and proteomes [15,16] (Fig 1A). To identify specific RBPs that are differentially regulated through temperature, we employed RNA Interactome Capture (RIC) [25] on A549 cells incubated for 48 h at 33°C or 37°C. This method allowed for the purification and identification of poly(A) RNA-specific RBPs via UV crosslinking and mass spectrometry (Fig 1B). Quality control of the protein samples to be analysed by MS was assayed by silver staining, highlighting the presence of distinct protein compositions between experimental conditions (Fig 1C). We observed clear differences in band patterns between the 33°C and 37°C in the pulldown samples indicative of the differences between the two proteomes, which were evaluated by LC-MS/MS. The resulting dataset was analysed for enrichment of RBPs in the 33°C compared to the 37°C samples (Fig 1D). Adjusted p-values were calculated with a cutoff of p<0.05. Of the total 423 RBPs identified across all datasets, 308 were found to be significantly regulated by alterations in growth temperature. Between these datasets, we identified a considerable number of RBPs that were preferentially enriched in either temperature. Interestingly, a total of 209 RBPs were shared between 33°C and 37°C (Fig 1E and 1F). Gene ontology analysis on the cold-responsive dataset, revealed a significant overrepresentation of cellular factors orchestrating mRNA stabilisation, nucleocytoplasmic export, and translational machinery (Fig S1). Interestingly, cellular component categorisation highlighted a pronounced enrichment of core components spanning distinct biomolecular condensates (nuclear specks, P-granules, and cytoplasmic stress granules), hinting at a coordinated, temperature-dependent spatial reorganisation of post-transcriptional ribonucleoprotein networks. Individual proteins that were present in both datasets, but significantly enriched at 33°C, included RBM3, YBOX1, HNRNPC, RBM47 and SND1 (Supplementary Table 1 and 2). Overall, these datasets provide important evidence that the RBP environment within a cell is extremely plastic, additionally highlighting the abundance of host factors that are selectively enriched at cold temperatures, that are possibly overlooked for their role in respiratory virus infections. Herein, we focused on the role of RBM3 in IAV replication, one of the top hits enriched in the 33°C dataset, as we previously identified this cold-inducible RBP with our collaborators through an IAV A/WSN/33 NP mRNA interactome screen [23].

**Figure 1.**
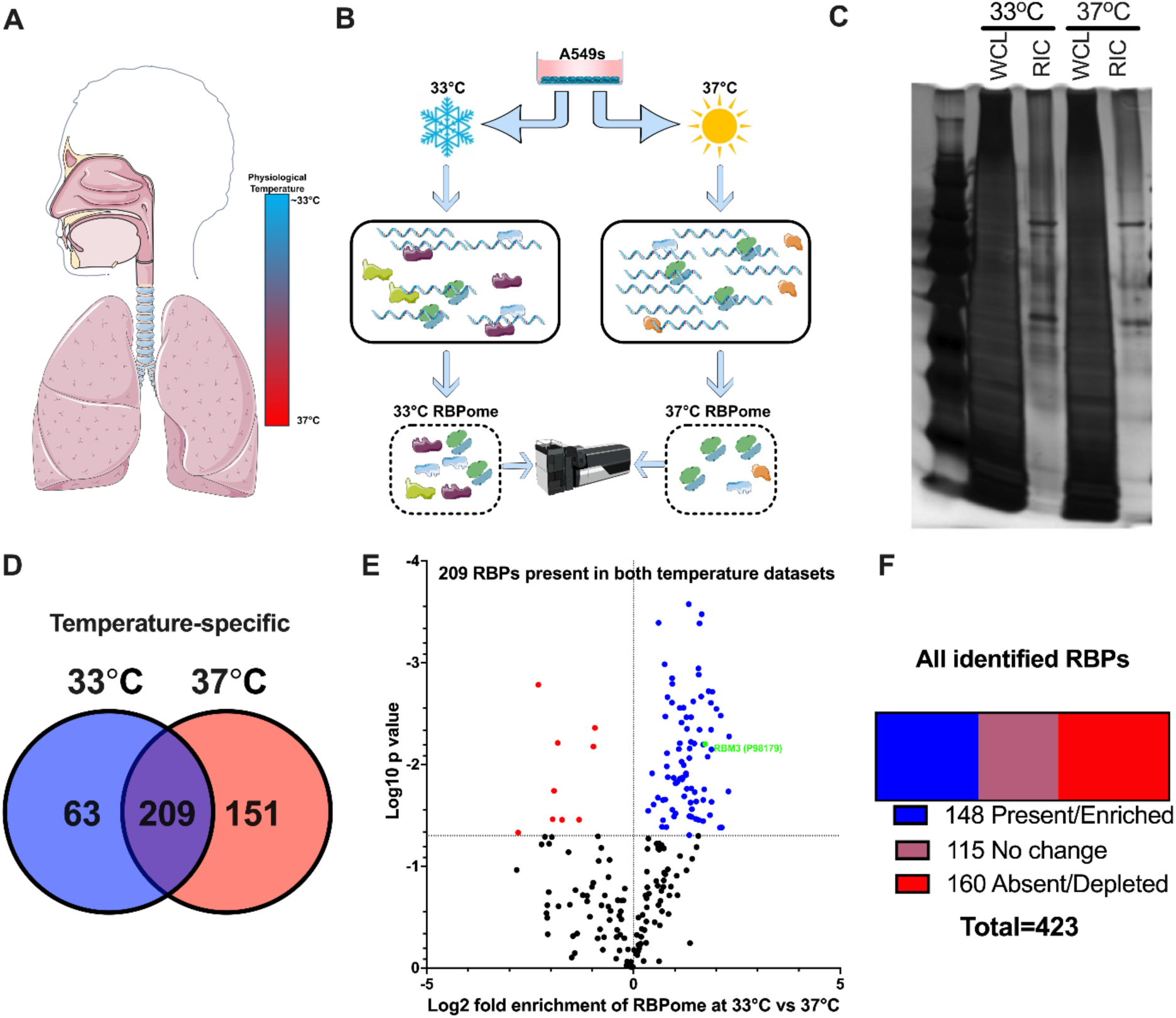
Cooler growth temperatures vastly alter the RBPome of A549 cells. **(A)** Schematic representation of the temperature gradient between the URT and LRT. **(B)** Schematic of the workflow for RIC in A549 cells incubated at 33°C or 37°C. **(C)** Representative silver stain of a gel with the RIC whole cell lysate input (WCL) and pulldown (RIC) samples from protein populations at 33°C and 37°C. **(D)** Venn diagram showing RBPs present in either 33°C datasets, 37°C datasets or both. **(E)** Volcano plot showing the enrichment of RBPs isolated from A549 33°C and 37°C pulldown samples. **(F)** Representation of the proteins significantly altered between datasets, whether by presence/absence or enrichment/depletion. Datasets derived from 3 independent biological samples.

### RBM3 is a positive regulator of IAV replication in A549s, mediated by its RNA-binding activity

In order to understand the effect of these two temperatures on virus replication, we incubated A549 cells at 37°C and 33° C for 48 h and analysed the replication kinetics of IAV A/WSN/33 (hereon referred to as WSN). As shown in Fig S2A the virus replicates significantly slower at 33°C when compared to virus replicating at 37°C. This was also confirmed by the delay in the production of viral mRNA levels from segments 5 & 6 (Fig S2B) in a single cycle replication at 33°C. Next, to investigate how RBM3 impacts IAV replication, we performed siRNA-mediated knockdown of RBM3 in A549 cells maintained at 37°C and 33°C, followed by infection for 8hpi (Fig 2A) and 16 hpi (Fig 2C) respectively. Titration of the virus containing supernatants revealed that knockdown of RBM3 results in a significant ∼2-log decrease in viral titres in both conditions. Knockdown efficiency of RBM3 was confirmed by Western blot (Fig 2B & 2D respectively). The decrease in viral titres upon RBM3 KD, correlates with the decrease in viral protein levels of nucleoprotein (NP) & polymerase basic protein 1 (PB1), indicating that RBM3 plays a proviral role in the virus life cycle before protein production at both LRT-& URT-like temperatures.

**Figure 2.**
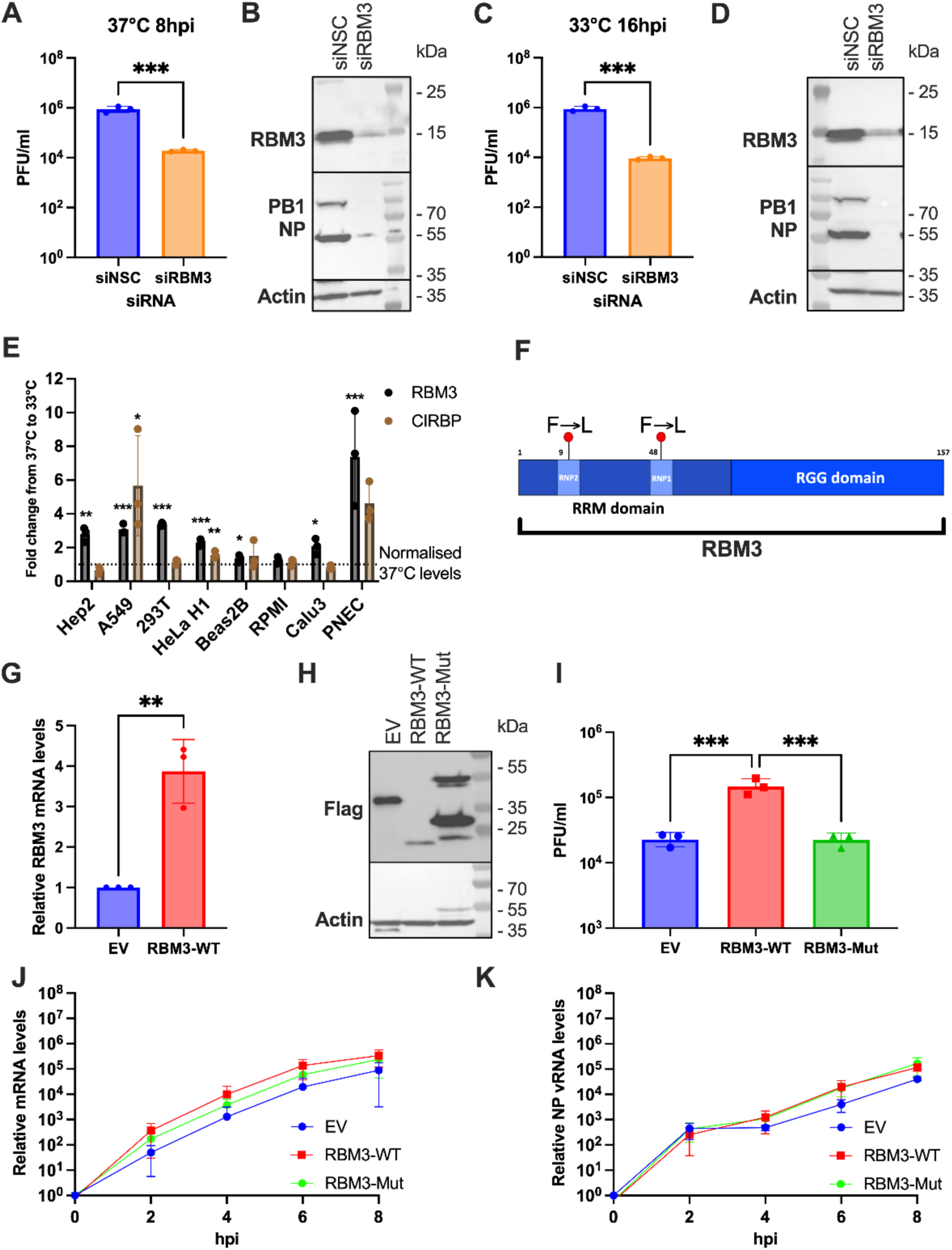
RBM3 possesses a proviral activity against influenza A viruses siRNA mediated knockdown of RBM3 was carried out in A549 cells maintained at 37°C. **(A)** and 33°C **(C)**, followed by infection with WSN (MOI 5). Virus titres were determined at 8hpi by standard plaque assays. A non-targeting siRNA was used as a negative control. Unpaired t-test of transformed data was performed to analyse significance with *** representing p<0.001. **(B)** & **(D)** The cells from (A) & (C) were lysed with laemmli buffer and levels of RBM3 and viral proteins (PB1 and NP) were analysed using SDS PAGE followed by Western blot; actin detection was used as loading control. Blots shown are representative of three independent experiments. **(E)** Cold responsiveness of a panel of immortalised cell lines and primary nasal epithelial cells (PNECs) as determined by the induction of RBM3/CIRBP expression by qPCR over 48 h and normalised to 37°C levels. Unpaired t-test of transformed data was performed to analyse significance with *, ** and *** representing p<0.1, p<0.01 and p<0.001 respectively. **(F)** Schematic representation of RBM3. Single nonsense mutations were introduced into the RNA binding domains, RNP2 and RNP1, creating an RNA-binding null mutant RBM3-Mut. **(G)** Levels of RBM3 mRNA in EV and RBM3-WT cellsgrown at 37°C were analysed by qPCR. Unpaired t-test was performed to analyse significance with ** representing p=0.0032. **(H)** EV, RBM3-WT and RBM3-Mut cells grown at 37°C were lysed with laemelli buffer and levels of Flag was analysed, with RBM3-Flag visible at ∼20 kDa, and secondary products arising in just the RBM3-Mut (potential dimers) appearing above this. Actin was used as a loading control. **(I)** EV, RBM3-WT and RBM3-Mut cells were grown at 37°C and infected with WSN (MOI 5) and 8 hpi, virus containing supernatants were titrated using standard plaque assay. One way ANOVA followed by Tukey’s multiple-comparisons test was performed to analyse significance with *** representing p <0.001. EV, RBM3-WT and RBM3-Mut cells were grown at 37°C and infected with WSN (MOI 5) followed by RNA extraction, cDNA synthesis and qPCR to analyse the levels of NP mRNA **(J)** & NP vRNA **(K)**. All data in graphs are shown as mean ± SD of three biological replicates.

Next, we sought to determine whether this proviral phenotype arises from a direct interaction between RBM3 and viral mRNA. However, as evidenced from the RIC-MS, the cold environment induces a global transcriptomic change, making it difficult to ascertain the effect of RBM3 alone on viral replication at 33°C. To circumvent this limitation, a cell line expressing cold-induced levels of RBM3 at 37°C was needed. First, a panel of cell lines commonly used to study respiratory viruses was assessed to determine their cold responsiveness, quantified by their ability to induce RBM3 when cultured at 33°C for 48 h. Primary nasal epithelial cells cultured at air-liquid interface (ALI-PNECs) were used as a positive control as it was expected that these would be highly responsive to cooler growth temperatures. RNA was extracted and the levels of RBM3 expression were quantified by qPCR, normalised to the levels observed with culturing at 37°C for each cell type (Fig 2E). Alongside RBM3, the transcript levels of another classical cold-inducible protein, CIRBP, were also quantified, with a similar pattern of cold-responsiveness observed across cell lines. While PNECs were the greatest responders, with an approximate 7-fold increase in RBM3 mRNA levels, A549s and 293Ts both demonstrated a >3-fold increase in RBM3 levels. Considering that immortalised cells can be manipulated to a greater extent than PNECs, we proceeded to use A549s to develop greater mechanistic insights into the role of RBM3 during IAV infection. Therefore, using a lentiviral system we generated a clonal A549 cell line that stably overexpressed the RBM3 CDS encoding for a C terminal Flag tag, termed RBM3-WT, that could then be cultured at 37°C. RBM3 is composed of two major domains; an RNA Recognition Motif (RRM) found at the C-terminus, and an N-terminus arginine-glycine rich (RGG) domain [26]. We hypothesised that if indeed RBM3’s proviral activity in IAV replication is dependent on its RNA-binding activity, an RNA-binding null mutant might serve as the perfect control. For this, we substituted two phenylalanine (F) residues to leucine (L) within the two highly conserved RNA binding regions RNP2 and RNP1 located within the RRM (Fig 2F) as this substitution was previously reported to disrupt RNA binding ability in other RBPs [27]. Similar to RBM3-WT, we also generated RBM3-Mut cell lines using the lentiviral system by transducing A549s with this RNA-binding null mutant encoding a Flag-tag. Additionally, a Flag-tagged Protein A expression vector was used to generate a cell line to serve as a negative control, termed EV. Overexpression of RBM3 in RBM3-WT cells was confirmed by qPCR, where transcript levels mimicked that of A549s cultured at 33°C (>3 fold over EV) (Fig 2G). Expression of these Flag-tagged proteins was validated by Western blot in all three cell lines (Fig 2H). To test if the proviral activity seen upon RBM3 KD is reproducible in these overexpression cell lines we infected EV, RBM3-WT and RBM3-Mut cell lines with WSN for 8 h, followed by titration of the virus-containing supernatant (Fig 2I). Accordingly, we observed a ∼log increase in titres in RBM3-WT cells when compared to EV, which in agreement with our hypothesis were abrogated to EV levels in RBM3-Mut cells, confirming that indeed RBM3’s proviral activity in IAV replication is mediated by its RNA binding activity. Since data from Fig 2B and D indicated to us that RBM3 plays a role in viral replication at a step preceding protein production, we tested if RBM3 impacts viral genome replication and transcription by analysing the levels of NP mRNA (Fig 2J) and NP vRNA (Fig 2K) in these three cell lines. Interestingly, we saw no significant changes in both these processes, suggesting that RBM3 potentially plays a role in the stages between transcription and protein production.

Recent work shows HNRNPH1 controls RBM3 via poison-exon splicing: inclusion of exon 3a in RBM3 pre-mRNA triggers nonsense mediated decay (NMD) [25], and weak HNRNPH1 (H1) binding at 37°C permits exon inclusion (low RBM3), whereas stronger binding at 30-33°C excludes exon 3a, boosting RBM3 levels [13]. To test this, we transfected A549s with siNSC or siH1 for 48 h and then treated with cycloheximide (CHX) or DMSO for 16 h and RNA was extracted followed by cDNA synthesis. Exclusion of exon 3a was analysed by agarose gel electrophoresis (Fig S3A) and quantified by performing RT-qPCR (Fig S3B). Both data revealed that exon 3a in only included in siH1+CHX samples confirming the H1 mediated RBM3 poison-exon splicing phenomenon in A549s. Knockdown of H1 also reduced RBM3 mRNA levels (Fig S3C). After WSN infection (MOI 0.01), siH1 cells had significantly lower NP mRNA at 48 hpi (Fig S3D) and viral titres were reduced to below limit of detection at 72 hpi (Fig S3E). Our data shows that HNRNPH1 mediated modulation of RBM3 expression by poison-exon regulation, interferes with IAV replication.

### RBM3 directly binds to influenza A virus transcripts with a preference for G-rich motifs within the NP mRNA

To determine whether the regulatory effects of RBM3 on IAV mRNA are mediated through direct physical interaction, we performed individual-nucleotide resolution UV cross-linking and immunoprecipitation (iCLIP2). Assays were conducted in our 3 A549 cell lines (EV, RBM3-WT, RBM3-Mut) undergoing a single round of viral replication and harvested at 6 h post-infection (hpi), a timepoint previously determined to coincide with maximal viral mRNA production in this cell line (Fig S2B). Characterisation was evaluated across three distinct conditions. Initial quantification of total viral reads per experiment demonstrated a robust enrichment of viral RNA capture in the RBM3-WT condition compared to both the EV and RBM3-Mut lines (Fig 3A), indicating that the interaction is specific and strictly dependent on the integrity of the RBM3 RNA-binding domain.

**Figure 3.**
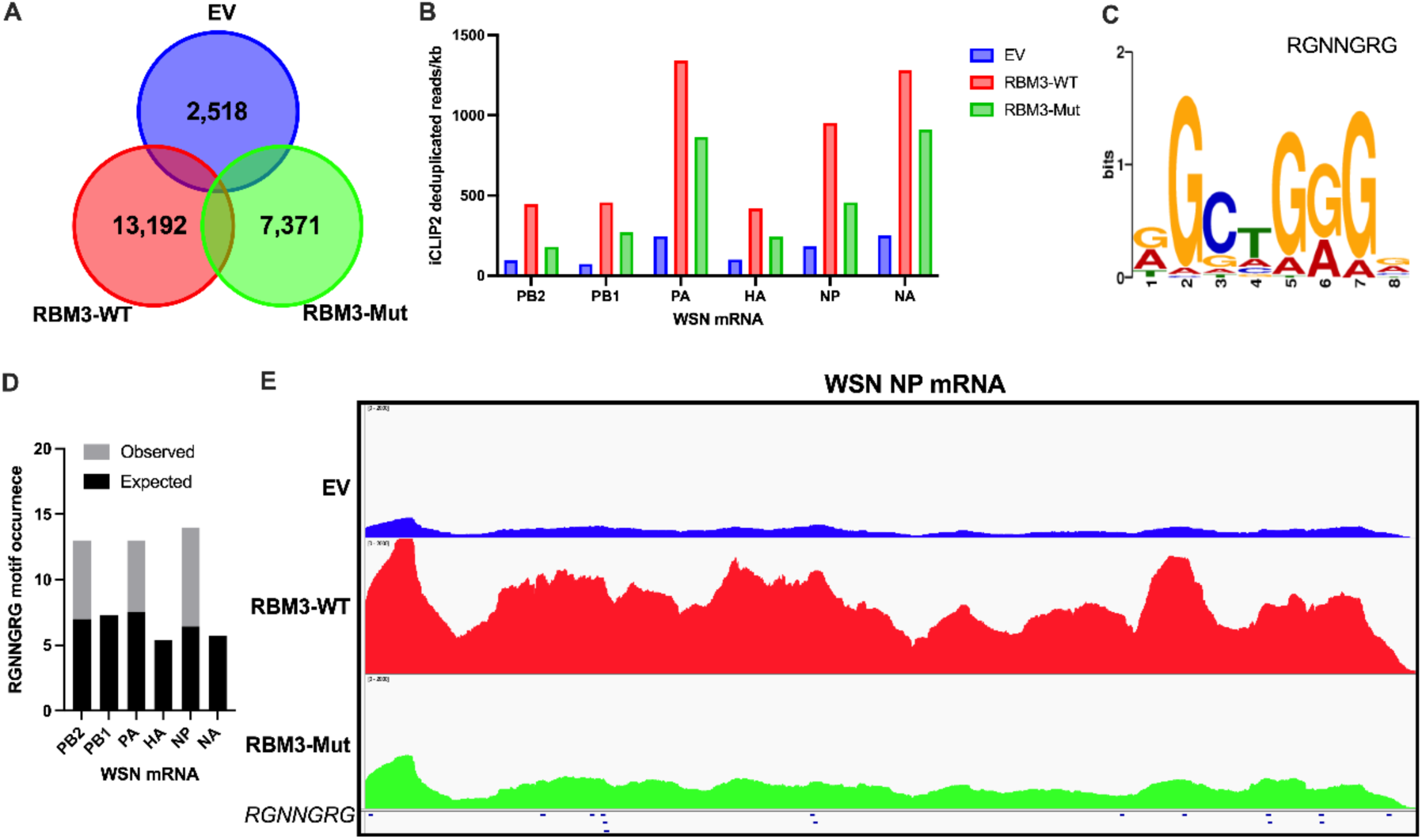
iCLIP2 maps direct RBM3 binding to influenza A virus transcripts via a specific G-rich motif during peak replication. **(A)** EV, RBM3-WT and RBM3-Mut cells maintained at 37°C were infected with WSN (MOI 3) to ensure a synchronized single round of replication and processed for iCLIP2 [27] at 6hpi to capture peak viral mRNA production. Venn diagram displaying the total number of viral cross-linked reads captured per experiment across EV, RBM3-WT and RBM3-Mut cell lines. **(B)** Quantification of iCLIP2 deduplicated read densities mapped across individual unspliced WSN viral mRNA segments (PB2, PB1, PA, HA, NP, and NA), normalised per kilobase (kb) of transcript for EV, RBM3-WT, and RBM3-Mut conditions. **(C)** Sequence logo of the highly enriched consensus motif (RGNNGRG) identified via de novo motif discovery using the MEME suite across high-confidence human RBM3-WT cross-linked interaction sites. **(D)** Combined distributional frequency bar chart of the RGNNGRG motif across individual unspliced WSN transcripts. Shaded bars indicate the analytical random background counts expected by chance, determined using a zero-order Markov baseline calculated from the exact mononucleotide composition of each separate viral segment. Overlaying grey sections indicate real observed motif frequencies, highlighting a distinct ∼2.2-fold enrichment within the target NP mRNA segment. **(E)** High-resolution Integrative Genomics Viewer track coverage maps showing the distribution of cross-linked iCLIP2 reads across the length of the unspliced WSN NP mRNA segment for EV, RBM3-WT, and RBM3-Mut expression lines. Mapped peak densities are shown relative to the precise spatial coordinates of predicted RGNNGRG consensus motifs, indicated by blue indicators along the lower track baseline.

To resolve the transcriptomic landscape of this interaction, we quantified the deduplicated reads mapping to individual unspliced viral mRNAs, normalised per kilobase of transcript (Fig 3B). While substantial binding was observed across multiple segments, notably PA, NP, and NA, the differential interaction between the WT and Mut proteins was highly non-uniform, indicating a specificity of RBM3 for some viral mRNAs over others. Intriguingly, the most pronounced effect was observed on the NP mRNA, which displayed an approximate 100% increase (a two-fold enrichment) in WT read density relative to the mutant control, marking it as a primary target for RBM3 coordination. This further supports our previous work that RBM3 is a direct interactor of NP mRNA using an RNA-interactome capture mass spectrometry approach [23].

To define the sequence determinants governing this recruitment, we subjected the cross-linked binding sites to de novo motif discovery using the MEME suite. This analysis identified a significantly enriched consensus motif, RGNNGRG (Fig 3C), consistent with the known preference of RBM3 for G-rich elements [26]. To rigorously evaluate whether this motif occurs more frequently than would be expected by chance across the viral genome, we established a segment-specific background baseline. Given the inherently skewed, Adenine-rich nature of the IAV genome, expected random counts for each segment were determined analytically using a zero-order Markov assumption based entirely on the exact mononucleotide composition of each individual unspliced A/WSN/1933 (WSN) transcript. Comparing these empirical baselines against real mapped occurrences via a combined bar graph revealed distinct target enrichment trends across the segments (Fig 3D). While segments such as PB1 (7 observed vs 7.32 expected) and HA (5 observed vs 5.38 expected) remained strictly at background levels, and NA showed distinct depletion (3 observed vs 5.75 expected), the NP transcript exhibited a striking ∼2.2-fold enrichment, yielding 14 real motifs against a background expectation of only 6.43 (Fig 3D). Notable elevations above background were also observed within another high-binding target, PA (13 observed vs 7.55 expected).

Finally, high-resolution visual inspection of the iCLIP2 binding profiles using the Integrative Genomics Viewer (IGV) confirmed that cross-linked reads spanned the length of the NP mRNA, with distinct, reproducible peaks directly aligning with the spatial positions of these predicted RGNNGRG motifs (Fig 3E). Taken together, these data demonstrate that RBM3 directly binds IAV transcripts during peak mRNA production, preferentially targeting specific G-rich motifs enriched within the NP mRNA.

### smiFISH-PLA confirms the direct interaction between RBM3 and NP mRNA

With data from Fig 2 indicating that RBM3 plays a proviral role at stages after replication but earlier than protein production and data from Fig 3 indicating the direct binding of RBM3 to NP mRNA, we next investigated if RBM3 affected the localisation of newly produced NP mRNA transcripts in these cells using our well-established smiFISH technique [17,28] (Fig 4A). Data from Fig S4C and S4D confirms the overall enhancement of RBM3 in the RBM3-WT cells when compared to both A549 as well as EV cells. Interestingly, we also observed the enhanced cytoplasmic localisation of Flag-tag in RBM3-Mut cells, in agreement with previously described observation that any mutation or truncation to the RRM/RGG region of RBM3, results in a predominantly cytoplasmic localisation [29]. This further indicated to us that the mutations we introduced in RBM3-Mut, indeed interfere with RBM3’s functions by mislocalising the protein. Colocalisation analysis between Flag and NP mRNA within the cytoplasm indicated a ∼4 fold increase in the Pearson’s correlation coefficient (PCC) in RBM3-WT cells over EV and ∼2 fold increase in PCC within the cytoplasm in RBM3-WT vs RBM3-Mut cells (Fig 4B). This data further confirmed that RBM3’s proviral role in the virus life cycle could potentially be physically interacting with NP mRNA. Since our EV cell lines were produced from lentiviral plasmids expressing a Flag-tagged Protein A, signal was detected in EV cell lines as well due to the direct binding of secondary antibody to the Protein A. Hence, we additionally imaged un-transduced A549s as a true negative control. With PCC values between Flag and NP protein averaging around 0 (1 indicates perfect positive colocalisation, 0 indicates no correlation and -1 indicates perfect negative correlation) in all cell lines, we were also able to rule out that RBM3 only interacts with NP mRNA transcripts and not the protein (Fig 4C). Interestingly, we saw significant enhancement in the localisation of NP mRNA within cytoplasmic region in both RBM3-WT and RBM3-Mut cells when compared to EV, but no significant difference when compared to A549 (Fig S4B). Whether this implies an increased trafficking of NP mRNA to the cytoplasm linked to regions of RBM3 RRM or RGG unmutated by us or simply a downregulation in EV cells due to the expression of Protein A needs further investigation.

**Figure 4.**
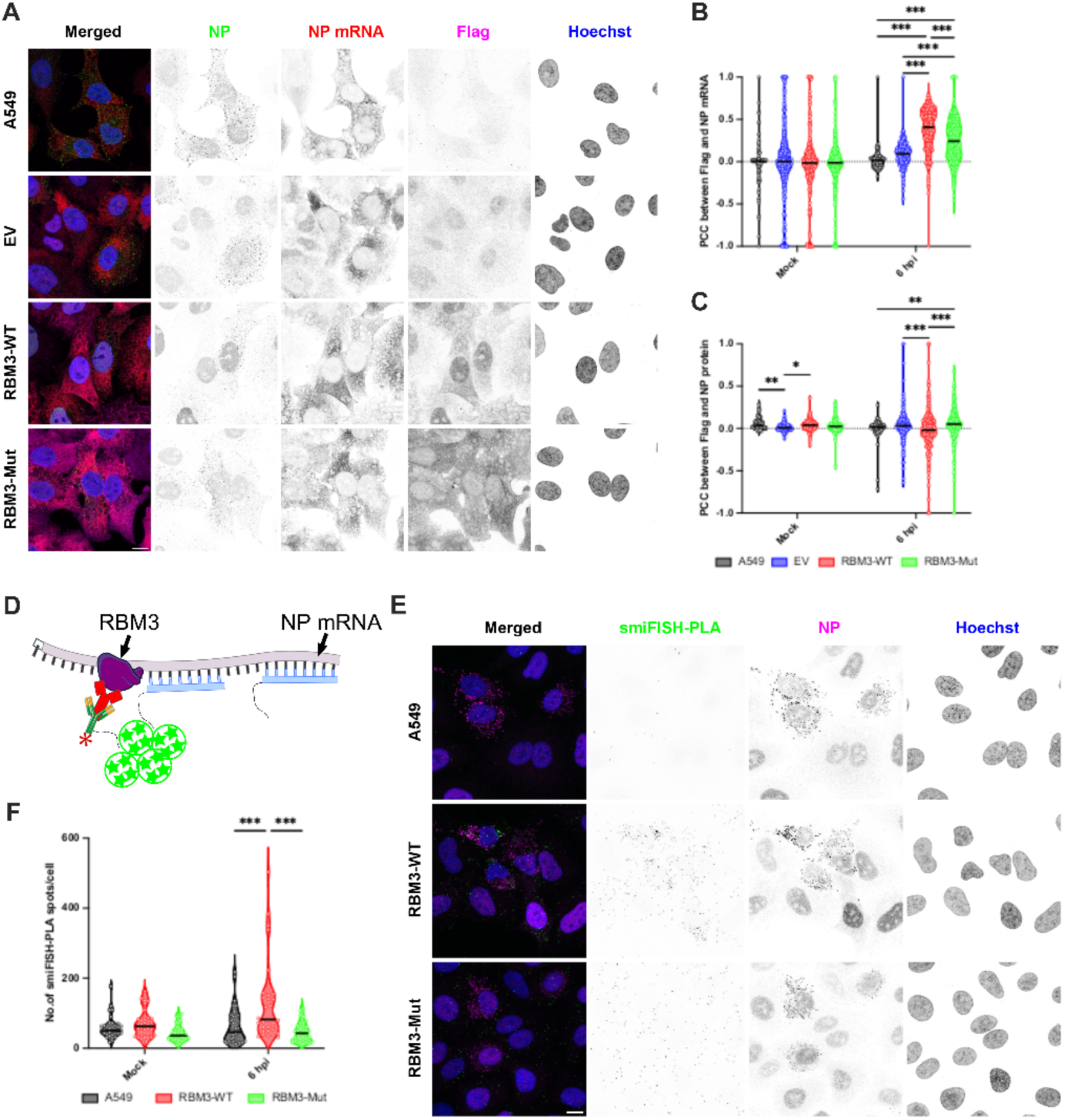
Direct interaction of RBM3 and NP mRNA visualised using smiFISH-PLA. **(A)** A549, EV, RBM3-WT and RBM3-Mut cells were maintained at 37°C infected with WSN (MOI 5), fixed at 6 hpi and processed for smiFISH to detect NP mRNA (Red) followed by Flag (Magenta) and IAV NP (green) detection using indirect immunofluorescence and counterstained with Hoechst. Pearson’s correlation coefficient quantification of the pairwise colocalisation between Flag and NP mRNA **(B)** as well as Flag and NP **(C)** in the cytoplasm was analysed in 87, 333, 344 and 281 cells across 3 biological replicates in these four cell lines. Two-way ANOVA followed by Tukey’s multiple comparisons test was performed to analyse significance with *, ** and *** representing p=0.04, p≤0.007. and p<0.001 respectively. Each point on the graph represents PCC values from a single cell. **(D)** Schematic of the smiFISH-PLA: NP mRNA was detected with smiFISH probes containing DuoLink PLA plus probes. Flag-tag was detected using a mouse anti-Flag primary antibody, followed by DuoLink anti-ms secondary antibody. smiFISH-PLA spots are subsequently generated by performing ligation and rolling circle amplification using the DuoLink PLA kit. **(E)** A549, RBM3-WT and RBM3-Mut cells were infected with WSN (MOI 5), fixed at 6 hpi and processed for smiFISH-PLA to detect the direct interaction between NP mRNA & RBM3 (Green) followed by IAV NP (Magenta) detection using indirect immunofluorescence and counterstained with Hoechst. **(F)** Number of smiFISH-PLA spots were quantified per cell. Two-way ANOVA followed by Tukey’s multiple comparisons test was performed to analyse significance with *** representing p<0.001. Each point in the graph represents the average number of smiFISH-PLA spots from 34 images from each cell line across three biological replicates. Scale Bar: 10 µM. Z stacks were acquired on Leica STELLARIS ONE 660 tauSTED Nanoscope with a 100x objective with 1.4 NA. Images presented are maximum intensity projections, processed with Fiji/ImageJ version 1.54p and analysed using CellProfiler version 4.2.6.

Since colocalisation analysis by PCC between two channels can often yield high correlation due to several factors including high protein concentrations, crowded intracellular environment and even the diffraction limit of light, we employed proximity ligation assay (PLA) to circumvent this limitation. Hence, to visualise true binding events between RBM3 and NP mRNA, we adapted a previously described RNA-PLA technique by combining it with our smiFISH protocol, referred here as smiFISH-PLA, the schematic of which is detailed in Fig 4D. [30]. Using this highly-novel, spatially resolved approach we were able to demonstrate for the first time where a direct interaction between a viral mRNA, NP, and a host RBP, RBM3, occurs intracellularly during infection (Fig 4E). Upon quantification, we found an ∼2-fold increase in number of smiFISH-PLA spots in RBM3-WT cells over both un-transduced A549 cells and the RNA-binding-deficient mutant RBM3-Mut (Fig 4F). These data not only indicated that RBM3 and NP mRNA are located less than 40nM apart but also validated that the mutations we introduced in the RBM3-Mut successfully abrogated the RNA binding activity of RBM3, once again confirming that RBM3’s proviral role in IAV replication is indeed mediated by its direct interaction with NP mRNA.

### Overexpression of RBM3 and subsequent binding to NP mRNA stabilises transcripts

Having visualised the interaction between RBM3 and NP mRNA, we next sought to identify how this interaction positively regulates WSN replication. Previous reports have suggested that RBM3 may stabilise certain bound transcripts in neuronal cells [21,22]. Given this context, we next explored the stability of a subset of viral transcripts in the RBM3-WT or EV cells. Cells were infected with WSN at an MOI 3 for 6 h, then treated with actinomycin D to prevent further global transcription [31]. Transcript levels were quantified hourly by qPCR over 5 timepoints from 0-4 h post-actinomycin D treatment to visualise the decay of RNA over this treatment period (Fig 5), with a one-phase decay model fitted to the data using a least squares regression. Observed differences in RNA stability were significantly different between cell lines and were assessed by comparing the decay constant (k) values using an extra sum-of-squares F-test, with p<0.05 deemed significant. Although we observed no significant effect on the stability of viral HA mRNA (Fig 5A), M2 mRNA did show a significant ∼50% increase in half-life (Fig 5B). However a robust and significant increase in stability was observed for NP mRNA (Fig 5C) with a >70% increase in RNA half-life. Actinomycin D assays performed in RBM3-WT or the RBM3-Mut cell lines revealed a loss of the stabilisation phenotype in the mutant cell line for viral NP transcripts (Fig 5D). These data suggest that RBM3 mediated stabilisation is a direct result of RBM3 binding to target transcripts, and that this is likely a form of cellular post-transcriptional regulation exploited by IAV during infection to promote viral transcript stability.

**Figure 5.**
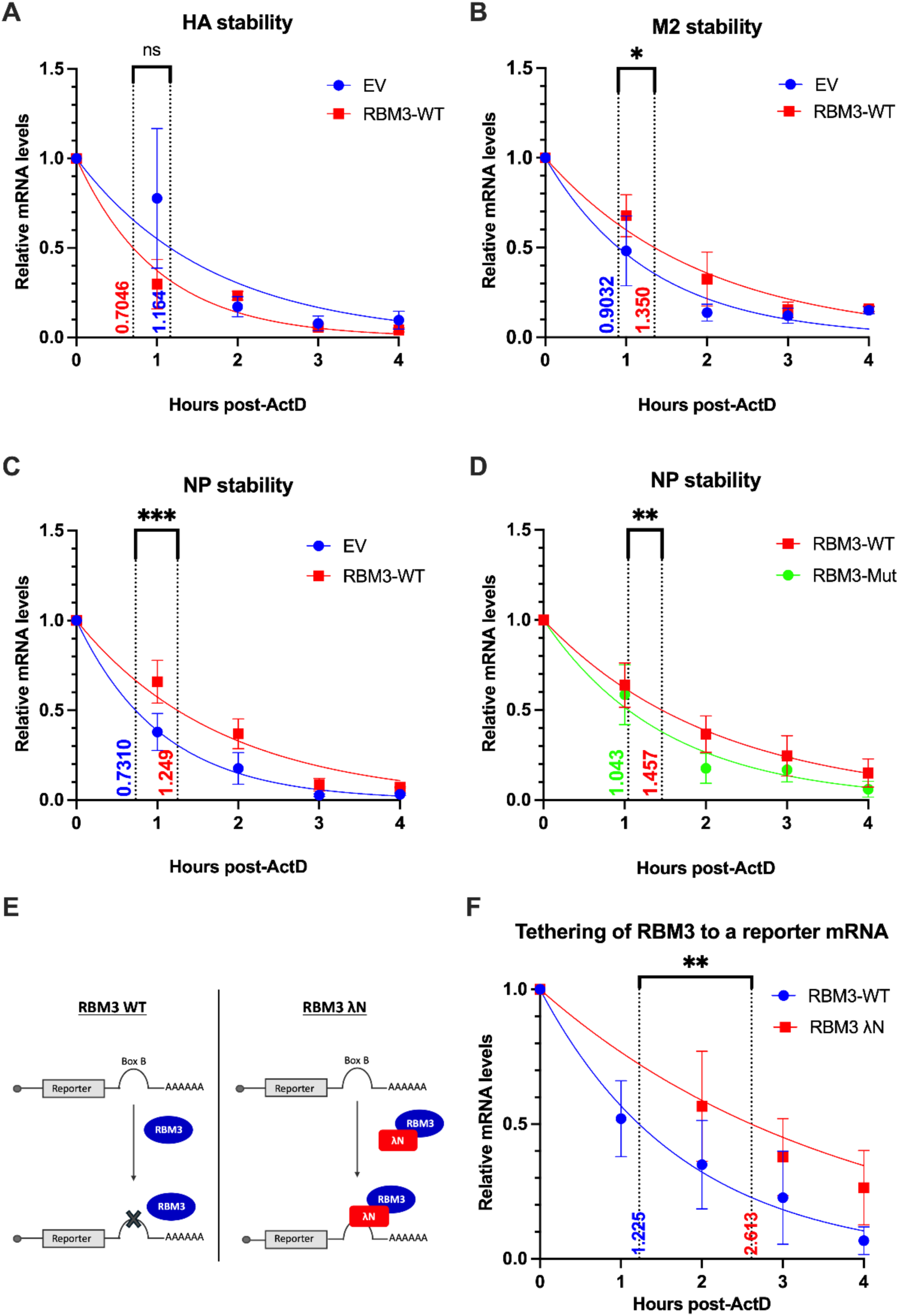
Direct binding of RBM3 to target mRNA including IAV NP mRNA enhances the transcript’s stabilisation. Transcript stability in EV or RBM3-WT cells grown at 37°C and was analysed by an actinomycin D (ActD) assay commenced at 6 hpi. Viral transcripts HA **(A)**, M2 **(B)** and NP **(C)** were quantified over 5 h. **(D)** The same assay was performed in RBM3-WT or RBM3-Mut cell lines to analyse NP mRNA decay kinetics. **(E)** Schematic representation of the tethering reporter assay. **(F)** 293T cells were transfected with a Luciferase BoxB reporter alongside either RBM3 or fused λN-RBM3. Cells were treated with Act D 48 h post-transfection. Luciferase transcripts were quantified over 4 h. Transcript half-lives in all the experiments were quantified by a one-phase decay assay using a least squares regression and differences in RNA stability were assessed by comparing the decay constant (k) values using an extra sum-of-squares F test, with *, **, *** representing p<0.05, p<0.01 and p<0.001 respectively. Data represented as mean ±SD of three biological replicates.

To be certain of our conclusions we employed a minimalist tethering approach. We set up a reporter system in which 293T cells were transfected with a Renilla Luciferase encoding for BoxB hairpins in the 3’ UTR, and either WT RBM3 or LambdaN (λN)-RBM3 fusion protein (Fig 4E). The 19 nt BoxB hairpin acts as the target sequence for λN binding and fusion of λN with an RNA-binding protein allows for elucidation of the RBP’s function on the BoxB-encoded reporter RNA [32]. Actinomycin D assays were repeated 48 h post-transfection and levels of luciferase RNA were assessed by qPCR. A significant enhancement in transcript half-life was observed upon expression of the fused λN-RBM3 protein (Fig 4F), providing further evidence that supports the role of RBM3 as an mRNA stabilising agent.

### Validation of RBM3’s proviral activity mediated by NP mRNA binding in primary nasal epithelial cells

Finally, we wanted to determine whether RBM3-mediated increase of overall viral replication would be mirrored in a clinically relevant model of PNECs. As previously shown in Fig 2E, PNECs were the cell lines that saw the highest increase in RBM3 levels when incubated at 33°C. In Fig 6A, similar to Fig S2A, we next investigated how UTR- and LTR-like temperatures affected IAV replication kinetics in monolayer cultures of PNECs (mPNECs). Interestingly viral titres were enhanced at 48 hpi by ∼1 log in PNECs grown at 33°C when compared to PNECs grown at 37°C. As nuclear-cytoplasmic shuttling of cold-inducible RBPs has previously been reported [33], we visualised the localisation of RBM3 in these mPNECs (Fig 6B). Not only were we able to visualise the overall increase of RBM3 expression in cells grown at 33°C quantified in Fig 6C as integrated intensity (total number of pixels) but also observed an increased cytoplasmic localisation when compared to the predominant nuclear localisation in mPNECs grown at 37°C. To confirm our findings from A549s that RBM3’s proviral role in IAV replication is due to its direct interaction with NP mRNA, we repeated our smiFISH-PLA protocol in mPNECs cultured at 33°C (Fig 6D). Progression of virus replication can be seen by the transition of predominantly nuclear localised of NP at 16 hpi to a more cytoplasmic distribution at 24 hpi (Fig S6A). Using this technique, we were able to confirm RBM3 and NP mRNA binding in PNECs as evidenced by the increase in the number of smiFISH-PLA spots over the course of the infection. To further validate the direct interaction between RBM3 and NP mRNA in these mPNECs we performed CLIP-qPCR in mPNECs. 24 hpi, UV crosslinking was performed and the lysates were subjected to lysis, immunoprecipitation of RBM3 protein, elution of bound RNAs and subsequent quantification by qPCR. NP mRNA was found to be greatly enriched in RBM3 pulldowns when compared to Actin mRNA level (Fig 6F). To further validate these mechanistic insights into RBM3 during IAV infection we proceeded to evaluate how altering RBM3 levels in highly clinically-relevant air-liquid interface PNEC (ALI-PNECs) cultures affects IAV replication. Previous studies have determined that targeted ASO treatment of the RBM3 poison exon, Ex3a in immortalised cell lines can result in the exclusion of the poison exon, preventing nonsense mediated decay (NMD) of RBM3 transcripts, and thus increasing overall RBM3 levels, effectively mimicking the function of HNRNPH1 [24]. Treatment of A549 cells with this previously published Ex3a-specific ASO for 48 h resulted in an almost 4-fold increase in RBM3 transcript levels compared to that of a non-targeting control (Fig 6G), confirming its effectiveness. Next, ASO treatment was applied apically to PNEC cultures from 3 donors. Primary cells are much less permissive to transfection therefore, ASO treatment would be predicted to be much less effective at inducing RBM3 in these cultures. Nevertheless, we observed a robust and significant ∼50% increase in RBM3 transcript levels after 48 h compared to the control ASO for all 3 donors (Fig 6H). Following this, we subjected PNEC cultures from the same 3 donors to ASO treatment followed by infection with WSN for 48 h. We then evaluated viral replication in these PNECS by quantification of viral RNA levels at 48 hpi. A significant increase in both NP mRNA and NP vRNA levels was observed by qPCR for each donor upon treatment with the Ex3a-specific ASO, with mRNA levels on average more than doubling after Ex3a ASO treatment (Fig 6I). These data confirm our earlier observations that increasing RBM3 levels positively regulates IAV replication, in a highly relevant human respiratory epithelial model of infection.

**Figure 6.**
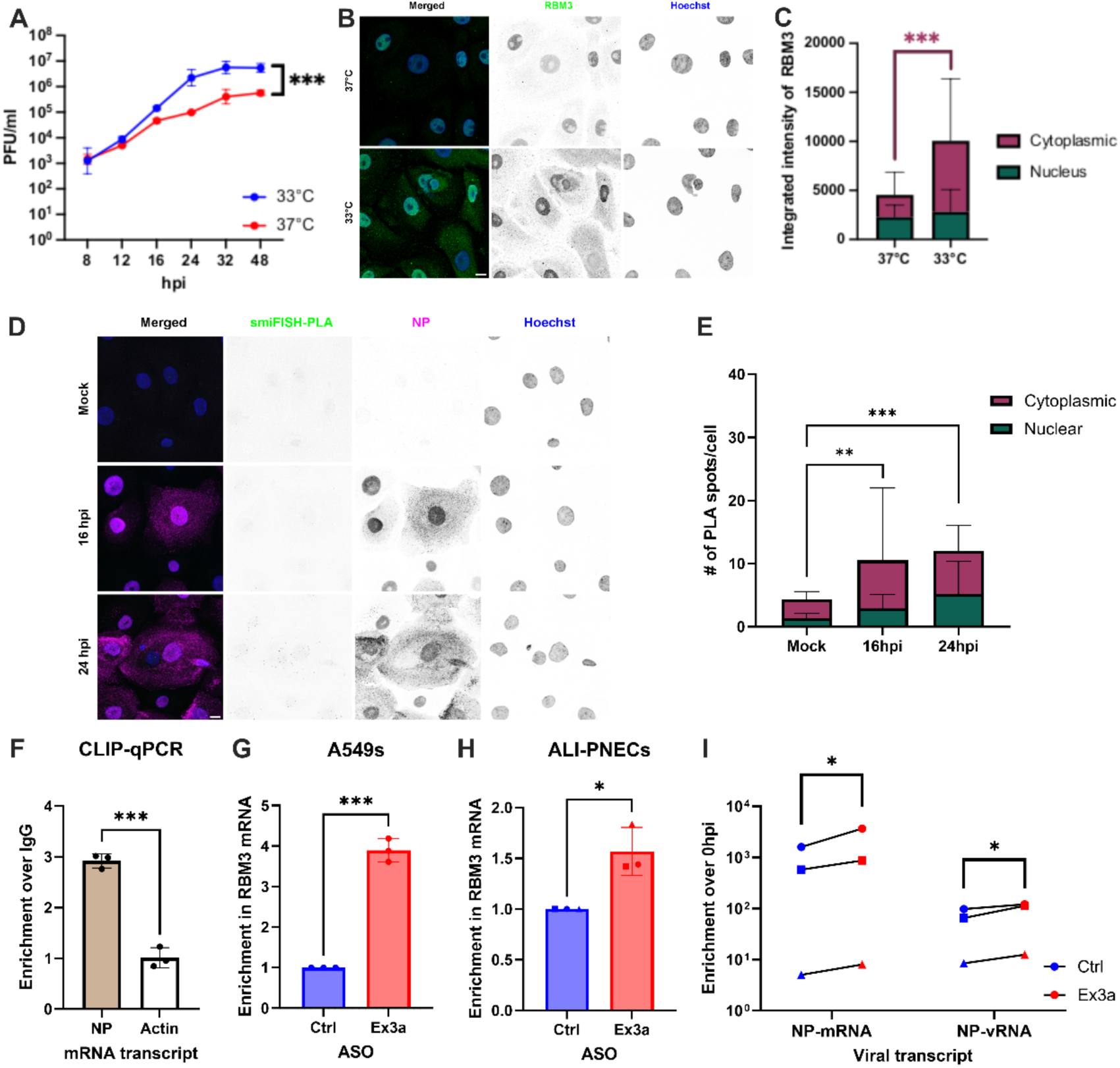
Validation of RBM3’s proviral activity in IAV replication mediated by its direct interaction with NPmRNA in primary nasal epithelial cells-. (A) Monolayer PNECS (mPNECS) were maintained at 37°C or 33°C and infected with WSN (MOI5). Virus titres were determined at the indicated time points by standard plaque assays. Two-way ANOVA followed by Tukey’s multiple comparisons test was performed to analyse significance with *** and **** indicating p.0.001 and p<0.0001 respectively. (B) mPNECs maintained at 37°C or 33°C were fixed and stained for RBM3 (Green) via indirect immunofluorescence followed by nuclear counterstaining with Höchst (Blue). (C) Total number of pixels (integrated intensity of RBM3) within nucleus and cytoplasm was calculated from 223 (33°C) and 291 (37°C) cells across three biological replicates. Two-way ANOVA followed by Tukey’s multiple comparisons test was performed to analyse significance with *** indicating p.0.001. (D) mPNECs maintained at 33°C were infected with MOI5 and fixed at indicated time intervals. Direct interaction of native RBM3 and NP mRNA was visualised using smiFISH-PLA followed by detection of IAV NP by indirect immunofluorescence and counterstaining with Höchst. (E) Average number of smiFISH-PLA spots in the cytoplasm and nucleus were quantified per cell from 219 (Mock), 199 (16 hpi) and 260 (24 hpi) cells from three biological replicates. Two-way ANOVA followed by Tukey’s multiple comparisons test was performed to analyse significance with ** and **** representing p<0.01 and p<0.0001 respectively. (F) mPNECs were maintained at 33°C and infected with WSN (MOI5), followed by UV-crosslinking at 24hpi and immunoprecipitation with anti-RBM3 antibody. qPCR quantification of CLIP samples demonstrated a significant enrichment in NP mRNA binding in comparison with Actin mRNA. Two-tailed unpaired t-test was performed with *** indicating p=0.0002. Data represented as mean ± SD of three biological replicates. (G) A549s maintained at 37°C were transfected with control (Ctrl) or Exon3a (Ex3a) specific ASOs. 48 h post transfection, RBM3 mRNA levels were determined using qPCR, with ∼4-fold increase. Two-tailed unpaired t-test was performed with *** indicating p<0.001. Data represented as mean ± SD of three biological replicates. (H) PNECs cultured at air-liquid interface (ALI-PNECs) from three different donors (indicated by different shaped data points), maintained at 37°C were transfected with ASOs and processed for qPCR similar to G. ∼50% ASO increase in RBM3 levels was observed with * indicating p=0.0143 after a Two-tailed unpaired t-test. (I) ALI-PNECs from three different donors (indicated by different shaped data points) were maintained at 37°C and transfected with Ctrl or Ex3a ASOs followed by infection with WSN (MOI 0.01). 48hpi, levels of NP mRNA and NP vRNA were analysed using qPCR. Two-tailed unpaired t-test revealed significance of * with p<0.05. All mPNEC and A549 data shown as mean ± SD of three biological replicates.

## Discussion

Given the continuing risk of respiratory virus pandemics, from coronaviruses such as SARS-CoV-2 or zoonotic influenza strains like H5N1, a detailed understanding of virus-host interactions in relevant tissue environments is critical for future preparedness. While highly pathogenic avian influenza strains have evolved to replicate at the gastro-intestinal tract of birds that exhibit ∼40°C to 42°C, influenza strains infecting humans have adapted to replicate efficiently in the upper respiratory tract (URT) that exhibits ∼33°C [34]. For a virus that transmits via aerosolised droplets, the nasopharyngeal URT is often the primary site of viral colonisation and a Glutamine to Lysine mutation in PB2 627 residue in the avian origin H5N1 strain has already been evidenced to aid in its adaptation to the cooler environment of the human URT [35]. Recent research from Turnbull et al., highlighted the importance in temperature differences during influenza replication showing that febrile temperatures could in itself pose as an antiviral defence mechanism [36]. However, how the cooler environment of the URT alters host cell biology and, in turn, aids in viral replication dynamics is currently unknown. We hypothesised that understanding how viruses exploit host factors in this unique tissue niche is essential to better comprehend transmission and avian influenza’s adaptation to the cooler temperatures of human URT. To this end, we present a comprehensive, temperature-resolved RBPome, identifying RNA-binding proteins (RBPs) that are differentially associated with mRNAs in A549 cells cultured at 33°C compared to 37°C (Fig 1). This is the first high-resolution characterisation of the cold-adapted RBPome in a context relevant to URT infection. We identified 63 RBPs unique to 33°C and 151 unique to 37°C, with 209 shared across both conditions. Among these, the cold-inducible protein RBM3 was significantly enriched at 33°C, suggesting temperature-dependent regulation of host-RNA interactions.

RBM3 is one of the evolutionarily conserved cold-inducible RBPs and is considered as a biosensor of cold temperature [37,38]. The activation of cold or hypoxic stress leads to the rapid upregulation of RBM3 transcription [33] and nucleocytoplasmic redistribution of the protein [13], where it binds target transcripts through its RNA-recognition motifs (RRMs) [33,39]. This binding then further promotes the stability of the target mRNA such as YAP1 and GAS6 [21,22], and thereby improving the translation efficiency of the stabilised transcripts such as COX-2, VEGF and IL-8 [40,41] and enhances cell-survival [39–41]. However, RBM3’s role in viral RNA regulation remains largely unexplored. Here, we show that RBM3 plays a proviral function during influenza replication by post-transcriptionally regulating viral mRNAs and enhancing their translational efficiency in the colder environment of URT by using an A549 overexpression cell line (RBM3-WT) and primary nasal epithelial cells (PNECs) as a URT model. Additionally, by introducing two substitutions in the RRM region of RBM3, we generated an RNA-binding null-mutant (RBM3-Mut) and were able to directly link RBM3’s RNA-binding activity to its proviral function.

Previous studies have utilised iCLIP2 to map RBP binding to viral RNA at high resolution during infections with HIV or SINV [42,43]. In this study, we utilised iCLIP2 for the first time in IAV infected cells and identified that amongst the eight IAV mRNAs, NP mRNA is the most enriched with the G-rich RBM3-binding motifs. Although this motif is similar to other G-rich motifs found on host RBM3-binding mRNAs, it is not entirely identical most likely due to differences in cell models and techniques used. Hettinger et al., performed RNA immunoprecipitation followed by RNA sequencing (RIP-seq) in C_2_C_12_ myotubes model [44] and Liu et al., used photoactivatable ribonucleoside enhanced crosslinking and immunoprecipitation (PAR-CLIP) in MEF cells [45], both identifying 3’UTR binding sites significantly enriched near polyadenylation sites. However, owing to the short length of 3’UTR sequence adjacent to the polyA tail of viral mRNA, we predict that the RNase I digestion, likely fragmented these 3’ UTR regions generating reads too small to be detected by iCLIP2. Additionally, we acknowledge the use of a single clonal A549 RBM3-overexpression line represents a potential limitation, as lentiviral integration may introduce unintended off-target effects. Despite this limitation, we were able to validate the enrichment of RBM3-binding motifs in NP mRNA with the identification of RBM3’s RNA-binding activity leading to the enhanced stabilisation of NP mRNA transcript but not HA mRNAs, a transcript that was found to have comparatively lesser RBM3-binding motifs from the iCLIP2 data.

We further employed multiple orthogonal assays to robustly validate this interaction. To directly visualise the intracellular localisation of RBM3 and NP mRNA interactions, we established smiFISH-PLA to detect close spatial proximity in situ (<40nM), a technique adapted from Alagia et al., [30] and found an enhanced interaction in the cytoplasm in RBM3-WT as well as PNECs cultured in monolayers. Previously truncation and mutations to RBM3’s RRM and/or RGG has been shown to affect its nuclear-cytoplasmic shuttling and thereby interfere with its anti-apoptotic functions [29]. Correspondingly, our data suggests that the integrity of the RRM region is essential to maintain the nuclear-cytoplasmic shuttling of RBM3 and thereby its interaction with viral NP mRNA. Our data collectively shows that any interference to the full-length protein (RBM3-Mut), disrupts its RNA binding activity and therefore, its interaction with NP mRNA and thereby, RBM3’s proviral functions during IAV replication. Our findings highlight the utility and robustness of smiFISH-PLA in mapping RBP-RNA interactions with subcellular resolution without the noise of biotin-labelling, even in clinically relevant primary cell models.

By using ASO-mediated induction of RBM3 [24], we further validate the translational relevance of our data by validating RBM3’s proviral role in a donor-specific manner. In agreement with our hypothesis, identifying the RBPome of the URT-like environment has led us to decipher the mechanism behind RBM3 mediated enhancement of viral replication in the URT, providing us with a potential target for future host directed antiviral strategies. Further investigation in appropriate animal models is required to understand if RBM3 also plays a role in transmission and aids in host-adaptation. While IAV appears to have evolved to exploit RBM3 to enhance replication efficiency in the URT, this mechanism could be repurposed therapeutically. Given the success of mRNA vaccines for SARS-CoV-2 [46], and ongoing development of nasal vaccine platforms to induce mucosal immunity [47,48], RBM3’s RNA-stabilising properties could be harnessed to enhance mRNA stability and translation in future respiratory therapies. Notably, recent advances have shown inhalable mRNA delivery to the airway epithelium *in vivo* [6], making this a viable direction for translational research.

In conclusion, RBM3 is a temperature-sensitive host factor that directly binds IAV mRNA post-transcription, regulating the transcript by stabilising it within the cytoplasm, enhancing protein production and thereby aiding in viral replication. Our multi-assay validation, the creation of a novel temperature-specific RBPome, establishment of smiFISH-PLA and the identification of a new functional axis in viral RNA biology collectively underscore the originality and impact of this work. Beyond IAV, our findings suggest that cold-inducible RBPs may represent a broader class of modulators shaping respiratory virus replication in epithelial tissues. These insights expand our understanding of host-pathogen interactions under physiologically relevant conditions and open new avenues for therapeutic intervention and pandemic preparedness.

## Methods

### Cells

All immortalised cell lines were cultured in DMEM supplemented with 1% Pen-Strep (Thermo Fisher Scientific; 15140122) and 10% FBS (Thermo Fisher Scientific; 10270106) at 37°C/33°C and 5% CO2. Cell lines used were A549s (ATCC; CCL-185), HEK 293Ts (ATCC; CRL-3216), MDCKs (CCL-34), Hep2s (ATCC; CCL-23), HeLa H1s (ATCC; CRL-1958), Beas2Bs (ATCC; CRL-3588), RPMI 2650s (ATCC; CCL-30) and Calu3s (ATCC; HTB-55).

### Monolayer and Well-Differentiated Primary Nasal Epithelial Cell Culture

Primary nasal epithelial cells were obtained from Promocell and monolayer (mPNECS) as well as well-differentiated primary nasal epithelial cells cultured in air liquid interface (ALI-PNECs) were established as described previously [49]. For monolayer cultures, PNECs were seeded directly onto 24 well dishes for Fig 6A and onto 24 well dishes with 12mm coverslips for imaging experiments in Promocell Airway Epithelial Cell Growth Medium Kit (C-21160) and were initially maintained at 37°C for two days to allow for cells to attach. Subsequently, cells were maintained at the indicated temperatures of 37°C or 33°C until confluency with medium changed every two days. For CLIP-qPCR, mPNECs were seeded in 10 cm dishes and maintained similarly at 33°C.

To generate ALI-PNECs, briefly, PNECs were seeded onto collagen coated 6 mm Transwells (Sarstedt, 83.3932.041) at 3×10^4^ cells/transwell. Cells were submerged in modified Promocell Airway Epithelial Cell Growth Medium which was replaced on the apical and basal side every 2 days until the cells reached confluency. Once confluent, an air liquid interface was initiated by completely removing the apical medium. Cells were differentiated using Pneumacult ALI medium (Stemcell Technologies, #05001) supplemented with hydrocortisone and heparin. For downstream experiments, ALI-PNECs were maintained at 37°C and only used when there was extensive cilia coverage and mucus production, clear hallmarks of full differentiation.

### Virus stocks and infections

IAV WSN (A/WSN/33) stocks were generated from a reverse genetics system previously described [50]. WSN stocks were grown on MDCK cells in IAV growth media consisting of DMEM, Pen-Strep, 0.2% BSA (Merck; A8412), 25mM HEPES (Merck; H0887) and 0.5 µg of TPCK-trypsin (Merck; T8802) and titrated on MDCK cells by standard plaque assay. The limit of detection (LOD) was defined as the lowest dilution at which plaques could be reliably distinguished and quantified by plaque assay, with this calculated to be 500 PFU/ml for all plaque assay titrations. A549s and MDCK IIs were infected with WSN in DMEM supplemented with 1% Pen-Strep (Thermo Fisher Scientific; 15140122) at the indicated MOIs for 1 hr at 37°C followed by removal of inoculum. The cells were then washed twice with PBS and incubated with infection medium (DMEM, supplemented with 1% Pen-Strep, 7.5% BSA and 1M HEPES) at the indicated temperatures. For multicycle replication kinetics in A549s, infection medium was supplemented with 0.2 µg of TPCK-trypsin.

#### RIC

The protocol employed here to capture the RNA interactome of poly(A)+ RNA (RIC) from A549 cells has been described previously [18,25]. At the time of seeding, 48 h before crosslinking and harvesting, A549 cells were either incubated under the normal growth temperature of 37°C, to determine the 37°C RBPome, or incubated in a 33°C incubator, to determine the 33°C RBPome. Cells were grown for 48 h at these conditions before performing UV crosslinking at 254 nm, 300 mJ/cm² and lysing cells in ice-cold PBS. The remaining steps were followed exactly as has been described previously [25].

#### SDS-PAGE and western blot analysis

Cells were washed twice with PBS and lysed in Laemmli buffer (1M Tris-HCl, pH6.8, 10% (w/v) SDS, 40% (v/v) glycerol, 0.01% (w/v) bromophenol blue) with 2% (v/v) β-Mercaptoethanol (Carl Roth) and sonicated for 30 min and denatured at 95°C for 10 min. Equal amounts of total protein were separated using Tris-glycine-SDS polyacrylamide gels (Invitrogen) and transferred onto nitrocellulose membranes (#1620112, BioRad). Total protein was detected by silver staining performed according to the manufacturer’s protocol, using the Pierce Silver Stain Kit (Thermo Fisher Scientific; 24612), for RIC samples. Protein detection was performed as has been previously described [51]. Primary antibodies for influenza PB1 (GTX125923, Genetex), NP (AB128193 (C43), abcam), RBM3 (Proteintech; 14363-1-AP), Actin (Proteintech; 66009-1-ig), and Flag (Proteintech; 66008-3-ig) were all used at a 1:5000 dilution.

### LC-MS/MS and data analysis

Mass spectrometry of RIC samples was performed at the Cambridge Centre for Proteomics. Sample treatment, preparation and trypsin digestion were performed as described previously [52]. Data for each sample were acquired on an Orbitrap Fusion Lumos Tribrid mass spectrometer over a 60 min run. No fractionation was performed due to the likely low complexity of the sample. Raw mass spectrometry data was analysed using MaxQuant software (v2.4.7.0) and utilising the Andromeda search engine [53]. The MS/MS spectra were aligned to the Uniprot Homo sapiens protein database, with known mass spectrometry contaminants and reversed sequences also included. The search was performed with trypsin selected as the specific enzyme and a maximum of 2 missed cleavage events allowed. The MS accuracy was set to 10 ppm, then 0.05 Da for the MS/MS. The maximum peptide charge was set to seven and seven amino acids were required for the minimum peptide length. One unique peptide to the protein group was required to call the presence of a specific protein, while a false-discovery rate (FDR) of 1% was set. Statistical analysis to determine enriched proteins in one set of biological samples over another was performed using the intensities quantified by MaxQuant on Perseus software (v2.0.11) [54]. First, any potential contaminants, reverse hits or proteins that were not well identified (fell below 1% FDR) were removed. Then only proteins that were identified in at least 2 replicates from a single biological condition were utilised. After filtering, the intensities were log transformed (log_2_) and statistical significance was calculated by way of an unpaired two-tailed Student’s t-test for each protein group. These values were then log transformed (-log_10_) and the resulting p values and enrichments were graphed as volcano plots. The cutoff for statistical significance was set to p < 0.05.

### Gene Ontology analysis

To functionally categorise the identified cold RBPome components, Gene Ontology (GO) term and protein class enrichment analyses were performed. The list of identified proteins was evaluated against the standard human proteome reference background using PANTHER. Enriched terms spanning Protein Classes, Cellular Components, and Biological Processes were filtered using a significance threshold determined by Benjamini-Hochberg False Discovery Rate (FDR) with a cut-off of *p* < 0.05. Fold enrichment was calculated as the ratio of the number of observed proteins in a given category to the expected number of proteins based on the reference proteome.

### siRNA knockdowns

A pair of siRNAs targeting the specified mRNA were used in each knockdown experiment, aside from when a single non-specific control siRNA (siNSC) was used. Sequences of these siRNAs are listed in Supplementary Table 3. siRBM3 and siH1 were used at a final concentration of 5nM and 20 nM respectively and transfected using Lipofectamine RNAiMAX (Invitrogen; 13778075) following the manufacturer’s instructions. A549 cells were reverse transfected and incubated for 24 h, after which media was changed with fresh growth media. Cells were then incubated for a further 24 h prior to infection.

### qPCR

RNA was extracted from cells using TRIzol (Invitrogen; 15596026). RNA was isolated and precipitated following the manufacturer’s instructions and 200 ng was reverse transcribed into cDNA using the ABI cDNA synthesis kit (Applied Biosystems; 4368814). For all cellular targets RT was performed using an oligo dT, while viral RNA underwent separate RT reactions with primers specific to each mRNA or vRNA [55]. Primers are listed in Supplementary Table 4 alongside the primers used for qPCR amplification. All qPCR experiments were performed in QuantStudio (Applied Biosystems). using SYBR Select Master Mix (Applied Biosystems; 4472908) following the manufacturer’s instructions. All qPCR data were quantified using the ΔΔ CT method.

### Establishment of a RBM3 overexpression cell line using a lentiviral vector

A lentiviral expression plasmid was generated encoding for either a full length wild-type RBM3 C-terminally tagged with 2X Flag epitope tags (RBM3-WT), a full-length RNA-binding null mutant (RBM3-Mut), containing two single phenylalanine to leucine substitutions within RNP1 and RNP2, or a Protein A expressing empty vector (EV). It should be noted that this RBM3 ORF did not contain the poison exon but instead encoded for just the open-reading frame. Protein A, RBM3-WT, and RBM3-Mut were followed by a P2A self-cleaving peptide and a puromycin resistance gene, similar to a system we have published previously [56]. Again, similar to previous publications, we rescued lentiviral particles by co-transfecting 293T cells with the RBM3-WT, RBM3-Mut or EV lentiviral vector alongside d8.74 (Addgene; 22036) and MD2G (Addgene; 12259). At 72 h post-transfection the supernatant was harvested, passed through a 0.45 μm filter and then overlaid onto fresh A549 cells. At 72 h post transduction cells were treated with 2 μg/ml puromycin and left to select for 72 h. Cells were then single cell cloned in 96 well plates in the presence of 2 μg/ml puromycin and left to grow for 3 weeks. Single cell colonies were then selected, expanded, and checked by Western blot to confirm the expression of the transgene.

### iCLIP2 data processing and motif analysis

Raw iCLIP2 reads were demultiplexed, and unique molecular identifiers (UMIs) were extracted prior to adapter trimming. Reads were aligned to a combined human and viral genome reference using Bowtie, and PCR duplicates were removed with UMI-tools. Deduplicated alignments were used directly for downstream analysis. For motif discovery, full-length sequences from aligned human reads were extracted and subsampled. Enriched sequence motifs were identified using the MEME Suite (v5.5.8) in DNA mode, searching for motifs of 6-15 nt in length. Motif logos were generated from the most significant MEME output.

### smiFISH & indirect immunofluorescence

A549s, EV, RBM3-WT and RBM3-mut cells were seeded in 24-well plates onto 12 mm coverslips. 24h post seeding, cells were infected with WSN at MOI 5 and fixed at 6 hpi with 4% PFA and permeabilised with 70% ethanol for 2 h at 4°C. Cells were briefly washed with FISH wash buffer (10% SSC, 10% formamide, 80% ddH2O) and incubated O/N at 37°C with smiFISH probes against WSN NP mRNA (Supplementary Table 6) annealed to a Cy3 imager strand, as we have previously described [17]. Next day, cells were washed twice with FISH wash buffer at 37°C for 30 min. Cells were stained with primary antibodies Rabbit anti-NP (PA5-32242, Invitrogen) and Mouse anti-Flag (66008-3-ig, Proteintech) to detect RBM3-WT/Mut for 1 hr at 37°C, followed by Goat anti-Ms AF647 (A32728, Invitrogen) and Goat anti-Rb AF488 (A11008, Invitrogen) secondary antibodies along with counterstaining for nucleus with Hoechst (ab228551, abcam) for 1 hr at 37°C.

PNECs maintained at 37°C or 33°C were fixed and processed for indirect immunofluorescence similarly allowing for 24 h of permeabilisation in ethanol at 4°C. Cells were blocked with 10% (v/v) BSA for 2 h at 37°C followed by primary Rabbit anti-RBM3 antibody (Proteintech; 14363-1-AP) for 4 h at 37°C, followed by Goat anti-Rb AF488 (A11008, Invitrogen) secondary antibody along with counterstaining for nucleus with Hoechst (ab228551, abcam) for 2 h at 37°C.

Coverslips were mounted using ProLong™ Diamond antifade mounting media (P36961, Invitrogen) and cured overnight at room temperature and imaged with a Leica STELLARIS ONE 660 tauSTED Nanoscope with a 100x objective with 1.4 NA. Z stacks were acquired and processed using Fiji/ImageJ version 1.54p.

### smiFISH-PLA

EV, RBM3-WT or RBM3-Mut were seeded on to a 12 mm coverslip in 24 well plates. 24 h later cells were infected WSN at an MOI of 5. The infection was allowed to follow through for 6 h to allow for peak viral mRNA production. 6 h later, cells were fixed with 4% PFA for 15 min at RT, then washed thrice with PBS. Cells were permeabilised with 0.5% Triton X for 10 min at RT, then washed thrice with PBS. Following a protocol adapted from Alagia et al. [30], coverslips were incubated with PLA-blocking buffer (2X SSC, 2% (v/v) BSA, 0.5% TritonX-100, 40 ng/µl of sssDNA and 40 ng/µl tRNA in DNAse free water) in a pre-warmed humidified chamber and incubated for 1 hr at 37°C. Cells were subsequently incubated with FISH-PLA probe buffer (2X SSC, 0.5% TritonX-100, 100nM smiFISH-PLA probes, 20 ng/µl of sssDNA and 20ng/µl tRNA in DNAse free water) in the humidified chamber in a hybridisation oven set at 70-80°C for 5 min (or 95C for 3 min), followed by 18 h incubation at 37°C. The next day, coverslips were placed into a 24 well plate and three washes were performed with 2× SSC wash buffer (2XSSC, 2% (v/v) BSA, 0.1% TritonX-100 in DNAse free water), followed by two washes with PBS at 20°C for 5 min with gentle rocking. Cells were incubated with Duolink Blocking Solution at 37°C for 1 hr, followed by primary antibody against Flag-tag (Sigma, F1804) diluted in Duolink Antibody Diluent (1:200) for 18 h at 4°C in a humidified chamber. The next day, coverslips were placed into a 24 well plate and three washes were performed with pre-warmed Wash Buffer A (0.88% (w/v) NaCl, 0.12% (w/v) Tris Base, 0.05% Tween 20, pH adjusted with HCl to 7.4) at RT for 5 min with gentle rocking. Cells were incubated with Minus PLA probes diluted 1:5 in Duolink Antibody Diluent in a humidity chamber at 37°C for 1 hr followed by three washes with Wash Buffer A. Duolink ligase buffer was diluted 1:5 in ddH2O. For 60 μL of ligase reaction solution, 1.5 μL of ligase enzyme was added to 58.5 μL of the diluted Duolink ligase buffer. Cells were incubated with this ligase buffer reaction solution in a humidity chamber at 37°C for 30 min followed by three washes with Wash Buffer A. Duolink amplification buffer was diluted 1:5 in ddH2O and protected from light. For a 60 μL of amplification reaction solution, 0.75 μL of polymerase enzyme was added to 59.25 μL of the diluted Duolink amplification buffer. Cells were incubated with this amplification reaction solution in a humidity chamber at 37°C for 1 hr and 40 min followed by two washes with pre- warmed Wash Buffer B (0.584% (w/v) NaCl, 0.424% (w/v) Tris Base,2.6% Tris HCl, pH adjusted with HCl to 7.5) at RT for 10 min with gentle rocking followed by a last wash with 0.01× Wash Buffer B. PLA signal was crosslinked with 2% PFA for 10 min at RT before proceeding to IF against viral NP protein with 1 hr incubation of primary (Rabbit anti-NP, PA5-32242, Invitrogen) and secondary (Donkey anti-Rabbit 647, A32795, Invitrogen) antibodies with Höchst to counterstain the nucleus at 37°C.

mPNECs were seeded on to a 12 mm coverslip in 24 well plates and grown at 33°C until confluency and infected with WSN at an MOI of 5. Cells were fixed at 16 and 24 hpi with 4% PFA for 15 min at RT, then washed thrice with PBS. Cells were permeabilised with 0.2% Triton X for 10 min at RT, then washed thrice with PBS. Similar to A549s, blocking was performed in a pre-warmed humidified chamber for 2 h at 37°C followed by PLA probe buffer hybridisation. Wash steps were carried out similar to A549s followed by blocking at 37°C for 2 h. Cells were incubated with primary antibody against RBM3 (Proteintech; 14363-1-AP) diluted in Duolink Antibody Diluent (1:200) for 18 h at 4°C in a humidified chamber. Washing steps with Wash Buffer A and secondary antibody incubations were performed exactly the same as above. Ligation reaction solution was supplemented with ATP (final concentration 1mM) and ligation reaction was carried out at 37°C for 30 min followed by three washes with Wash Buffer A. Amplification step was performed for 2h and 30 min followed by two washes with pre-warmed Wash Buffer B at RT for 10 min with gentle rocking and a last wash with 0.01× Wash Buffer B. PLA signal was crosslinked with 2% PFA for 10 min at RT before proceeding to IF for NP with 2 h of incubation for primary (Rabbit anti-NP, PA5-32242, Invitrogen) and secondary (Donkey anti-Rabbit 647, A32795, Invitrogen) antibodies with Höchst to counterstain the nucleus at 37°C.

Coverslips were mounted on to slides with Diamond antifade Mounting medium and Z-stack images were acquired on Leica STELLARIS ONE 660 tauSTED Nanoscope with a 100x objective with 1.4 NA.

#### Image Analysis

Images presented are a maximum intensity projection of Z-stacks, processed using Fiji/ImageJ version 1.54p [57] with final image panels generated using the ImageJ plugin ScientiFig [58]. Scale bar 10 µM. Image analysis for nuclear-cytoplasmic segmentation, pixel quantification, smiFISH-PLA spots measurement and Pearson’s Correlation Coefficiency (PCC) were carried out using CellProfiler version 4.2.6 [59].

### Actinomycin D assays

RBM3-WT, RBM3-Mut and EV cells were seeded one day prior to infection in 24 well plates. Cells were infected with WSN at MOI 3. At 6 hpi, RNA was harvested from a single well for each cell type, constituting the 0 h timepoint. All remaining infected wells underwent media exchange for growth media containing 2 µg/mL Actinomycin D (SBR00013). RNA was harvested hourly from 7-10 hpi representing 1-4 h post Actinomycin D treatment. RNA underwent cDNA conversion using oligo dT or WSN specific primers, with the ABI cDNA synthesis kit (Applied Biosystems; 4368814). Quantification of the cDNAs was performed by qPCR as described above. Fold enrichment of RNA levels was quantified using the ΔΔCT method and normalised to either A549-EV or RBM3-WT levels.

### λN tethering assay

λN was cloned into a lentiviral expression vector expressing full-length RBM3. 293T cells were seeded in a 12-well plate the day before transfection. The following day, cells were transfected with 500 ng of pCI-RLuc-BoxB[60] and 100 ng of either the WT RBM3 or the λN-RBM3 WT expressing plasmids. At 48 h post-transfection, RNA was harvested for each transfection to represent a 0 h timepoint. All remaining cells were treated with 2 µg/mL Actinomycin D (SBR00013). RNA was harvested by TRIzol hourly up to 4 h post-treatment. RNA underwent DNaseI (Invitrogen, EN0521) digestion to remove any residual plasmid DNA before cDNA conversion using oligo dT primers with the ABI cDNA synthesis kit (Applied Biosystems; 4368814). Quantification of the cDNAs was performed by qPCR, described above. Fold enrichment of luciferase RNA levels was quantified using the ΔΔCT method and normalised to the corresponding 0 h mRNA levels.

### CLIP-qPCR

The CLIP-qPCR protocol employed here has been published previously [61]. Briefly, mPNECs were seeded in 10cm dishes and grown at 33°C as mentioned above. Cells were infected with WSN at MOI 5. At 24 hpi media was removed from the cells, before being washed in ice cold PBS. UV crosslinking was performed at 254 nm. Cells were then scraped, pelleted and lysed in 250 µl of NP40 lysis buffer (HEPES-KOH pH 7.5 50 mM, KCl 150 mM, EDTA pH 8.0 2 mM, NaF 1 mM and NP40 0.5%) on ice for 30 min. Lysates were clarified by centrifugation and 10% of the sample was saved as an input and boiled in Laemmli buffer containing 2% β-mercaptoethanol at 95°C for 10 min for western blotting. The remaining 90% of lysate underwent immunoprecipitation with anti-RBM3 antibody (Proteintech; 14363-1-AP) or IgG control antibody coupled to Protein G Dynabeads (ThermoFisher, 10004D) for 1 h at room temperature. Beads were then captured, washed five times with 1 mL NP40 lysis buffer and three times with PBS. A fraction (10%) of the beads was removed, resuspended in Laemmli buffer and boiled at 95°C for 10 min for Western blotting. The remaining 90% of beads underwent proteinase K digestion (ThermoFisher, EO0491) at 55°C for 60 min to release bound RNA, following the manufacturer’s instructions. RNA was then isolated from the digestion by TRIzol LS (ThermoFisher, 10296028) and ethanol precipitation and cDNA were synthesized using oligo dT or A/WSN/33 specific primers [55] (Supplementary Table 4). Quantification of the cDNAs was performed by qPCR, as described below. Fold enrichment of RNA levels was quantified using the ΔΔCT method and normalised to input levels.

### ASO transfection of A549s and ALI-PNECs

A549s were transfected with control or Exon 3a targeting ASOs [24] (Supplementary Table 5). ASOs were resuspended in DMEM at a concentration of 1 µM and transfected using Lipofectamine 2000 (Invitrogen, 1 µL/mL). At 48 h post-transfection RNA was harvested, cDNA synthesised as described above, and RBM3 levels quantified by qPCR. For ALI-PNECs, the transfection conditions were the same, with the ASO-containing transfection mix added apically. Cells were incubated for 48 h prior to infection with WSN MOI 0.01. At 1 hpi, the inoculum was removed and transwells were washed 5 times in PBS to remove unbound virus. RNA was harvested from WD-PNECS using Trizol™ either before infection or at 72 h post-infection.

### Statistical analysis

Unless otherwise stated, all experiments were performed as 3 independent biological replicates. When bar graphs are used to represent the data, each individual replicate can be observed as a point overlaid on the bar, with error bars indicating standard deviation. Significance was calculated as described in each figure legend with * indicating p<0.05, ** indicating p<0.01 and *** indicating p<0.001.

## Acknowledgements

This research was funded in part by an ERC-STG grant, PTFLU 949506 awarded to D.G.C. This research received infrastructure support from the Wellcome-Wolfson Institute for Experimental Medicine at Queen’s University Belfast. Authors would like to thank Dr Seyed Mehdi Jafarnejad for providing the plasmids for the BoxB reporter system.

In Figure 1 cells bilayer icon by Marcel Tisch is licensed under CC0, upper-respiratory-tract icon by Servier https://smart.servier.com/ is licensed under CC-BY 3.0 Unported, lung icon by Servier https://smart.servier.com/ is licensed under CC-BY 3.0 Unported, all protein icons by Servier https://smart.servier.com/ is licensed under CC-BY 3.0 Unported, Liquid_Chromatograph_Mass_Spectrometer icon by DBCLS https://togotv.dbcls.jp/en/pics.html is licensed under CC-BY 4.0 Unported and RNA icon by Servier https://smart.servier.com/ is licensed under CC-BY 3.0 Unported. In Figure 3 RNA, oligo, RBP and antibody icons were acquired from the NIAID NIH BIOART under a CC-BY license.

## Supplementary data

Supplementary Table 1 - Initial RIC dataset prior to downstream analysis

Supplementary Table 2 - RIC dataset of RBP identified in at least 2 replicates of each condition, with fold enrichment and p values included.

Supplementary Table 3 - siRNA sequences

Supplementary Table 4 - Primer sequences

Supplementary Table 5 - Sequences of antisense oligonucleotides

Supplementary Table 6 - smiFISH probe sequences

**Supplementary Figure 1.**
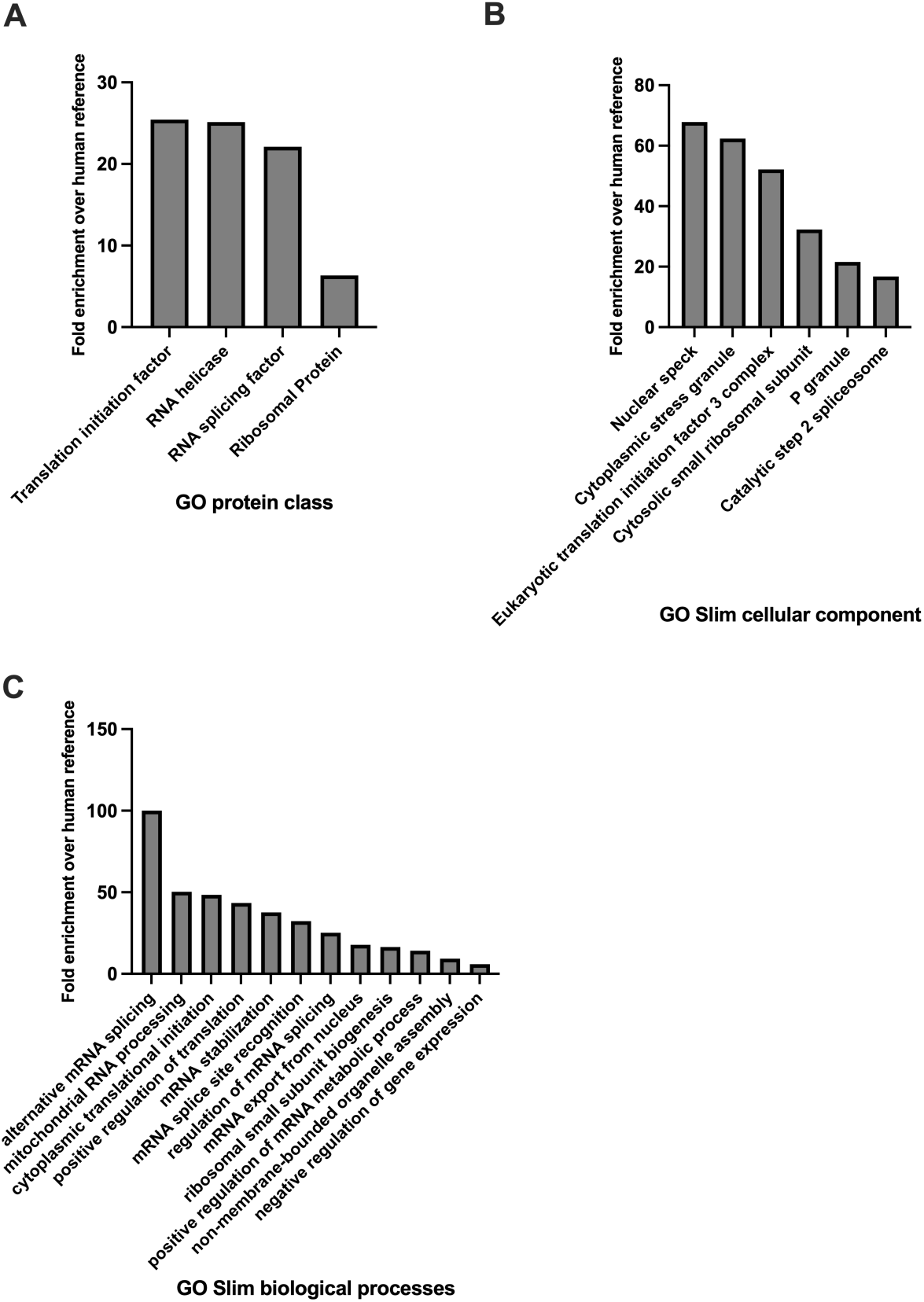
Functional characterisation and Gene Ontology enrichment analysis of the cold RBPome. **(A)** Enriched PANTHER protein classes mapped within the cold RBPome dataset. **(B)** Enriched GO Cellular Components highlighting subcellular localization. All data are presented as fold enrichment over the human reference proteome. **(C)** Enriched GO Biological Processes associated with identified proteins. Statistical significance was determined using Benjamini-Hochberg False Discovery Rate correction with only terms achieving p < 0.05 displayed.

**Supplementary Figure 2.**
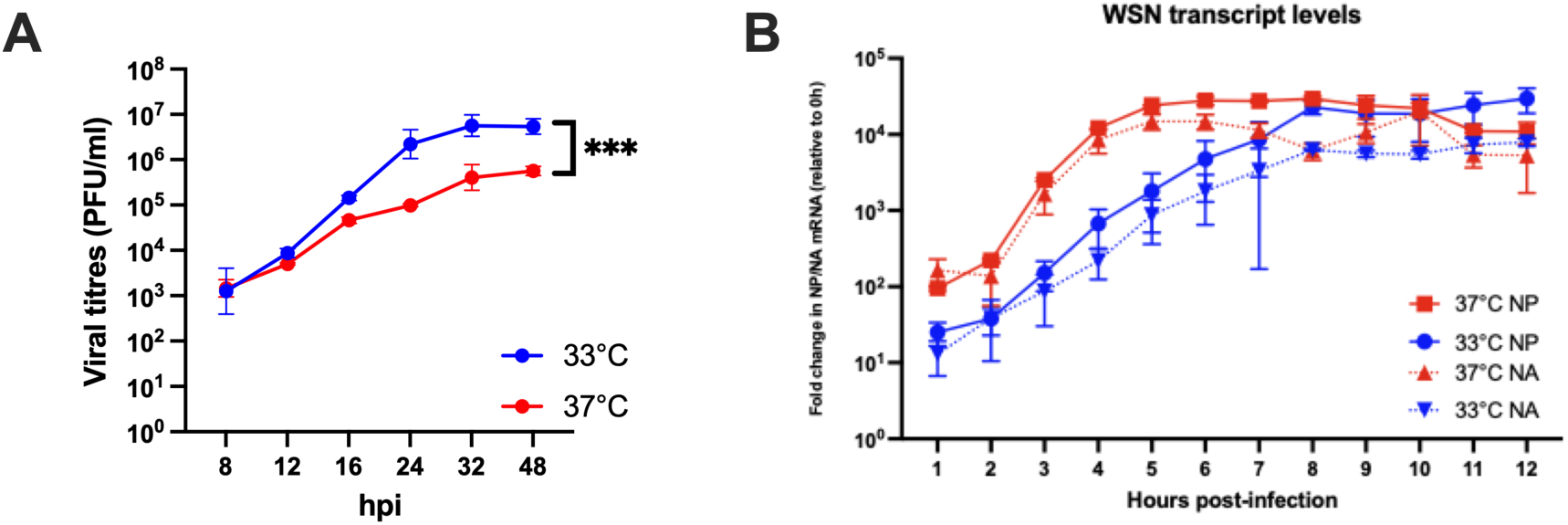
Reduction of RBM3 levels through manipulation of HNRNPH1 expression suppresses IAV replication. **(A)** A549 cells maintained at 37°C and 33°C were infected with WSN (MOI 0.01). Virus containing supernatants were harvested at the indicated timepoints and titres were quantified by standard plaque assay. Area under the curve is plotted with significance of *** indicating p <0.001. Data represented as mean ± SD of three biological replicates. **(B)** A549s maintained at 37°C and 33°C were infected with WSN (MOI 5). Cells were lysed for RNA extraction every hour from 0 hpi until 12 hpi and levels of NP & NA mRNA were analysed by qPCR. Data plotted as fold change of each transcript over their corresponding 0 hpi.

**Supplementary Figure 3.**
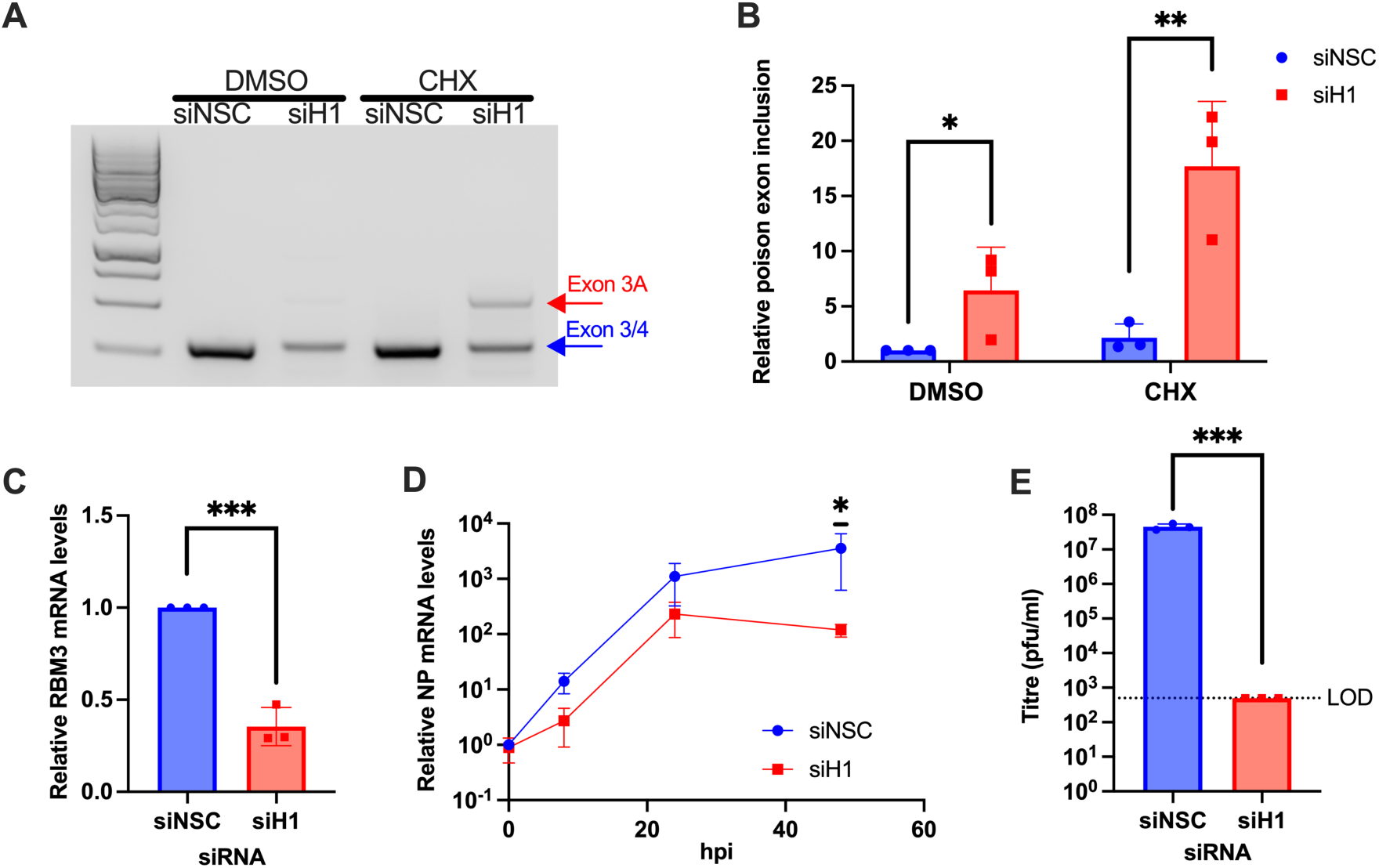
Reduction of RBM3 levels through manipulation of HNRNPH1 expression suppresses IAV replication. (A) A549s maintained at 37°C were transfected with HNRNPH1 siRNA (siH1). 48 hours post transfection cells were treated with DMSO or cycloheximide (CHX) for 16 h followed by RNA extraction. Inclusion of RBM3’s poison exon, exon 3a was observed by agarose gel following PCR. Representative blot of three biological replicates shown. (B) cDNA from S3A was further analysed by qPCR to quantify relative RBM3 poison exon 3a inclusion. Student’s T-test was performed to calculate significance between siNSC and siH1 treated cells, *=p<0.05, **=p<0.01. Data represented as mean ± SD of three biological replicates. (C) A549s maintained at 37°C were transfected with siH1. 48 hours post transfection, RBM3 mRNA levels were analysed by qPCR. Area under the curve is plotted with significance of ** indicating p <0.01 (D) A549s maintained at 37°C were transfected with siH1. 48 hours post transfection, cells were infected with WSN (MOI 0.01), and cells were lysed for RNA extraction at the indicated time points. qPCR analysis followed by statistical analysis by Student’s T-test revealed a significant difference in NP mRNA levels at 48hpi with * indicating p <0.05 (E) Virus containing supernatants from A549s transfected with siH1 (37°C) were harvested at 72 hpi (WSN, MOI 0.01) and quantified by standard plaque assay. Limit of detection (LOD) indicated by black dotted line (500 PFU/ml). All data shown as mean ± SD of three independent experiments.

**Supplementary Figure 4.**
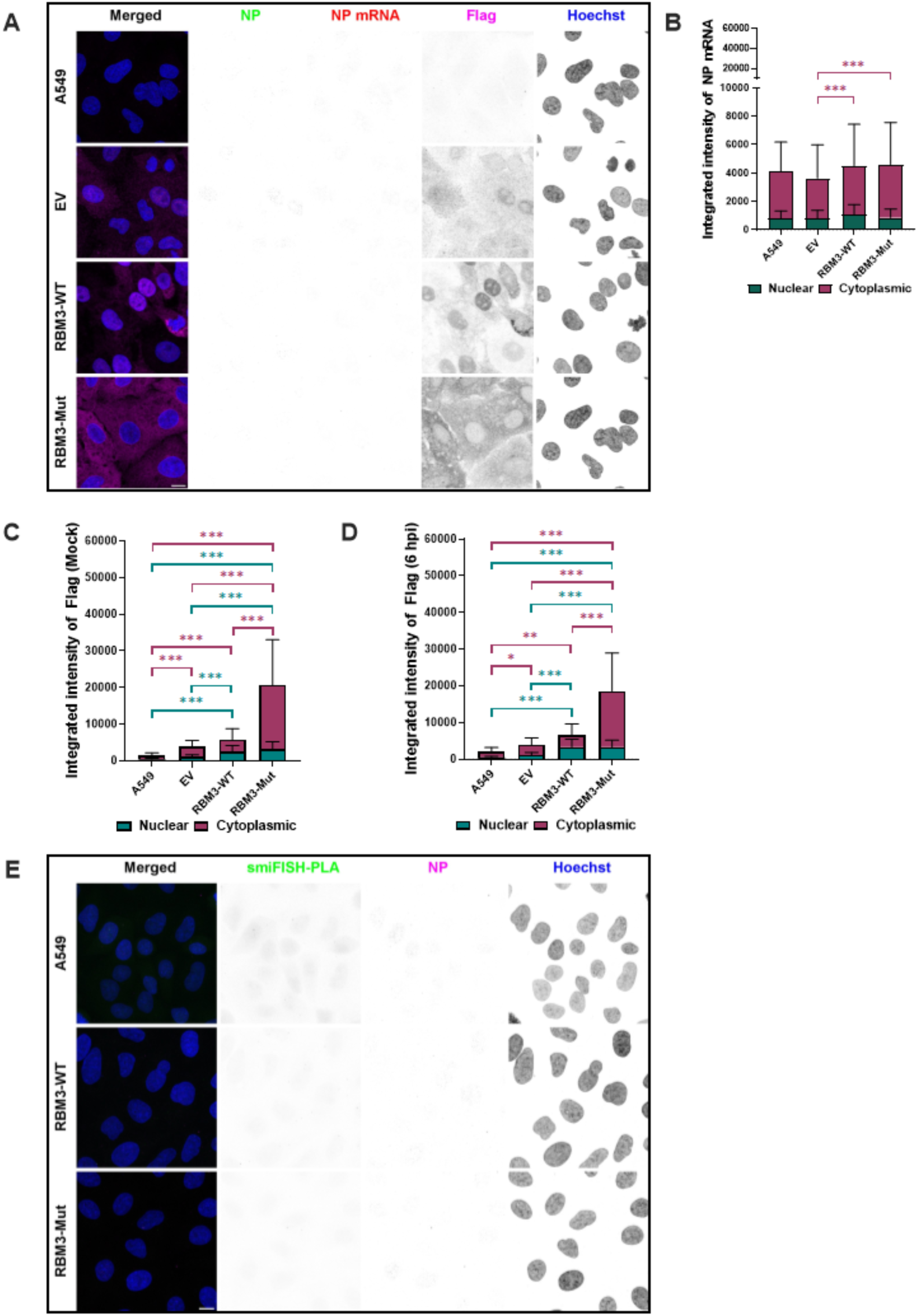
Immunofluorescence and smiFISH-PLA in mock infected A549s. **(A)** Mock-infected A549, EV, RBM3-WT and RBM3-Mut cells corresponding to Fig 4A. NP mRNA (Red), Flag (Magenta), IAV NP (Green) and Höchst (Blue). Total number of pixels (integrated intensity) of **(B)** NP mRNA 6 hpi **(C)** Flag Mock **(D)** Flag 6 hpi was calculated within nuclear and cytoplasmic region from 88 (A549), 333 (EV), 345 (RBM3-WT) and 279 (RBM3-Mut) cells across three biological replicates corresponding to Fig 4A. Two-way ANOVA followed by Tukey’s multiple comparisons test was performed to analyse significance with *, ** and *** representing p<0.05, p< 0.002 and p<0.0001 respectively. **(E)** smiFISH-PLA spots from mock infected A549, RBM3-WT and RBM3-Mut cells corresponding to Fig 4E. smiFISH-PLA spots (Green), IAV NP (Magenta) and Höchst (Blue). Scale Bar: 10 µM. Z stacks were acquired on Leica STELLARIS ONE 660 tauSTED Nanoscope with a 100x objective with 1.4 NA. Images presented are maximum intensity projections, processed with Fiji/ImageJ version 1.54p and analysed using CellProfiler version 4.2.6

